# Entorhinal and ventromedial prefrontal cortices abstract and generalise the structure of reinforcement learning problems

**DOI:** 10.1101/827253

**Authors:** Alon B Baram, Timothy H Muller, Hamed Nili, Mona Garvert, Timothy E J Behrens

## Abstract

Knowledge of the structure of a problem, such as relationships between stimuli, enables rapid learning and flexible inference. Humans and other animals can abstract this structural knowledge and generalise it to solve new problems. For example, in spatial reasoning, shortest-path inferences are immediate in new environments. Spatial structural transfer is mediated by grid cells in entorhinal and (in humans) medial prefrontal cortices, which maintain their structure across different environments. Here, using fMRI, we show that entorhinal and ventromedial prefrontal cortex (vmPFC) representations perform a much broader role in generalising the structure of problems. We introduce a task-remapping paradigm, where subjects solve multiple reinforcement learning (RL) problems differing in structural or sensory properties. We show that, as with space, entorhinal representations are preserved across different RL problems only if task structure is preserved. In vmPFC, representations of standard RL signals such as prediction error also vary as a function of task structure.

## Introduction

Reinforcement learning (RL) theory has given deep insights into the brain’s algorithms for learning, but has remained relatively mute about the representations that are the foundation for this learning. For example, how might a task, or an element of a task, be represented in the brain? Some recent progress has been made through comparison with spatial navigation, where representations are better understood. It has been suggested that the same representations that map Euclidean space (such as hippocampal place cells and entorhinal grid cells) may be extended to a broad range of non-spatial problems. In these cases, instead of representing physical location, they may represent location in an abstract space that captures the regularities of the task at hand^1–8^.

One attractive corollary of these ideas is that non-spatial tasks might benefit from the profound representational efficiencies that are known in space. In space, entorhinal cortex (EC) grid cells encode unique locations in the most efficient manner possible, given the correlation structure of observations on a 2D Euclidean plane^4,9–11^. Furthermore, in remapping experiments, this representational structure is transferred across environments^12–14^. Hence, in new spatial environments, it is not necessary to re-learn the associations implied by the structure of 2D space. Instead it is sufficient to learn what sensory observation is where in the map 7 and inferences (such as shortest paths^15,16^) can be made immediately. The ability to transfer structural knowledge in non-spatial tasks would, similarly, bestow efficiencies. The structure of a problem, learnt in one situation, could be mapped onto a new situation with different sensory observations, and solutions could immediately be inferred. Behaviourally, it is clear that both *humans* and animals profit from such efficiencies. In psychology, this phenomenon is known as “learning-set”^17^ - subjects with prior exposure to the structure of a problem are routinely better at solving new examples.

Might similar mechanisms support both spatial and non-spatial generalisation? Importantly, this would not require Euclidean-specific properties of grid cells to underlie non-spatial inferences. The hexagonal pattern, its phase, orientation, and scale all have no obvious parallels in abstract non-Euclidean RL tasks. Instead the neural computations that are supported by these representations may be maintained across Euclidean and non-Euclidean domains -s for example, the ability to encode the relationships between different states of a task, and generalise them to new problems. Indeed, these same computational principles *can* lead to efficient inferences in Euclidean and non-Euclidean environments, but do so using different underlying representations^4,6,7^.

For this to be true, one prerequisite is that brain regions that contain structural representations in Euclidean spatial tasks (like grid cells) should also represent the structure of a non-Euclidean RL task. A second is that, like grid cells in Euclidean spaces, these representations should (a) generalise across different sensory exemplars of the same structural problem and (b) differ between two problems of different structures.

In humans, grid-like coding has not only been recorded in EC, as in rodents, but also in a network of brain regions in association cortex^18–21^ including ventromedial prefrontal cortex (vmPFC). Notably, these brain regions are often also implicated in RL tasks^22^, where signals commonly reflect algorithmic variables such as value^23^ and prediction error^23–25^. If these variables have different consequences in different task structures, they may also conform to the representational predictions above^5^. Instead of a unitary representation of prediction error, these regions might generalise prediction error representations across different problems with the same task structure, but have different representations across problems with different structures.

To answer these questions, we designed an RL task where we manipulated either the problem structure (by changing the correlation structure of serial bandits) or the sensory stimuli in a 2×2 factorial design. This design mirrors a spatial remapping experiment with two important differences: (1) It is non-spatial task elements that are being remapped, as opposed to locations in 2D space. (2) We include 2 separate structures as well as 2 separate environments (sensory stimuli) in each structure. This design therefore enabled us to test for representations of the task structure, factorised from the representations of the stimuli they were tied to.

Using fMRI in humans, we found that the EC contained a representation that differed between different task structures, but generalised over different sensory examples of the same structure. Similarly, learning signals in regions including vmPFC and ventral striatum maintained different voxelwise patterns for different task structures, again generalising over different environments with the same structure.

## Results

### Task

Subjects performed a task where three 1-armed bandits were interleaved pseudo-randomly. Two of the bandits (bandits A & B) had correlated outcome probabilities, while the third (C) was independent. Crucially, we manipulated two features of the task across the different blocks: 1) the sign of the correlation between the A & B bandit probabilities. We refer to this correlation as the relational structure of the stimuli. 2) The stimuli set, with two possible triplets of images. Thus, there were 4 block-types, each with a specific combination of a relational structure and a stimuli set, arranged in a 2×2 factorial design (Figure 1b). The fMRI experiment comprised of 8 blocks of 30 trials each (10 trials per stimulus), divided into 2 independent runs of the 4 block-types, with a pseudo-random block order counterbalanced across subjects. Hence, in total subjects completed 8 blocks, 2 of each block-type.

In each trial, subjects viewed one of the three stimuli and had to indicate their prediction for its associated binary outcome (a “good” or a “bad” outcome, demarked by a King of Hearts or Two of Spades card, respectively) by either accepting or rejecting the stimulus (Figure 1a and S1a). Thus, there was always one correct answer in each trial: subjects should accept a stimulus if they predict the outcome to be the “good” outcome, and should reject if they predict the outcome to be the “bad” outcome (Figure 1a). Only in accepted trials, the subject would either win or lose a point, depending on the outcome. Outcome identity was revealed in all trials, including rejection trails, even though the subject’s score did not change in these trials (Figure 1a). Predictions of the outcomes could be formed based on the recent history of outcomes. The outcome probabilities switched pseudo-randomly between 0.9 and 0.1 with an average switch probability of 0.15. As the two correlated bandits switched together (Figure 1c), subjects could use their knowledge of the correlation structure to learn from the outcome on one related stimulus about the other.

The outcome probabilities associated with the related stimuli were negatively correlated (-Corr pairs) in half of the blocks (Figure 1 c, left two panels), and positively correlated (+Corr pairs) in the other half (Figure 1c, right two panels). In all blocks the third stimulus had an outcome probability which was uncorrelated with the other two stimuli (0Corr pairs). The current block-type was signalled by the background colour of all stimuli in the block. Subjects learned the mapping between background colour and correlation structure prior to scanning. Consequently, the only learning performed during scanning was of outcome probabilities, not of the relational structure - knowledge of which was available from the first scanning trial.

**Figure 1.**
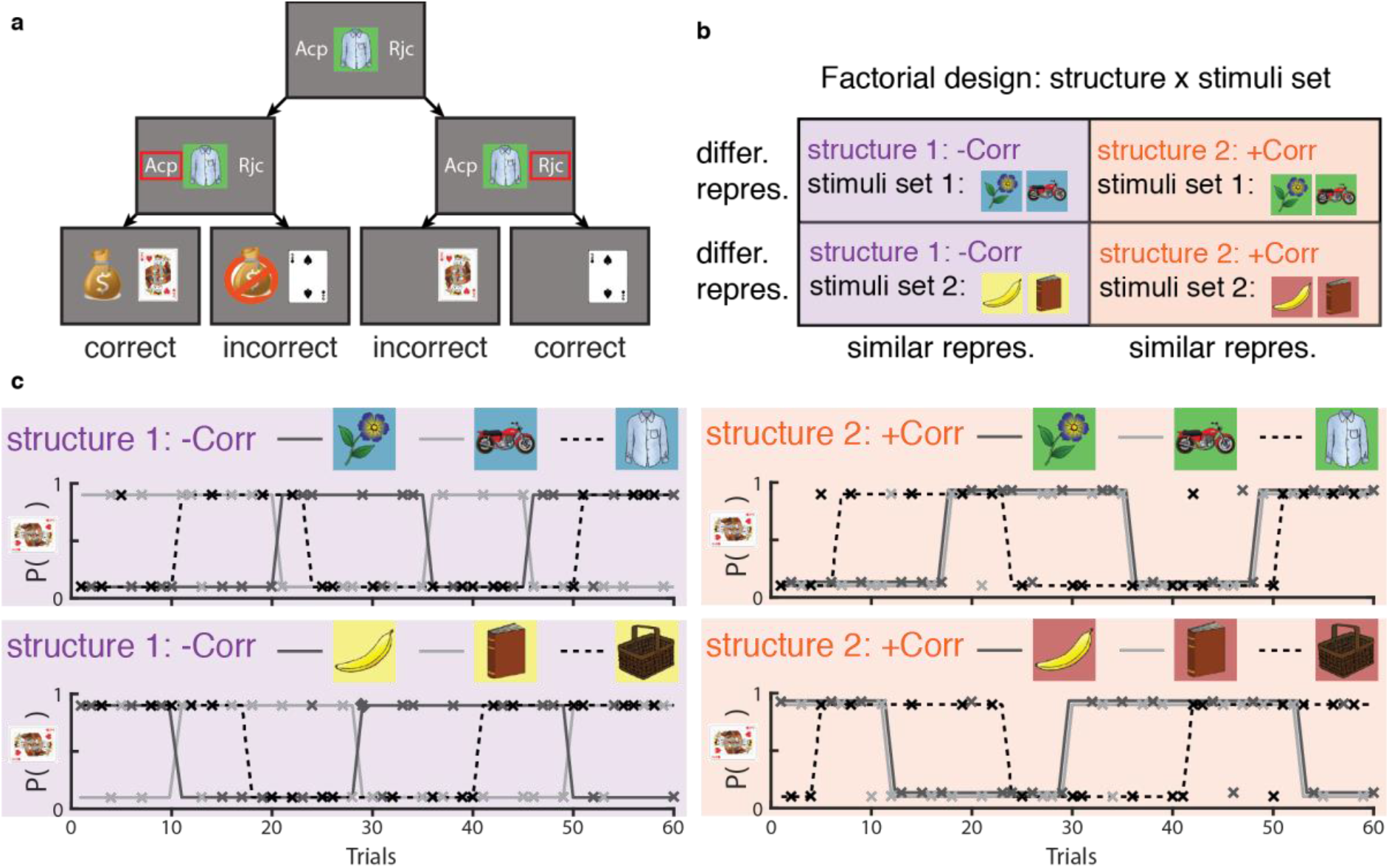
Task design. **a.** Possible progressions of a single trial. **b.** Experimental design and neural predictions for structure-encoding brain regions: 2×2 factorial design of stimuli set x relational structure. **c.** Example of the reward schedule for one subject in the four block-types. Solid grey lines and dashed black line are the probabilities of a good outcome for the related stimuli and the control stimulus, respectively. Xs mark the stimuli (colour) and actual binary outcomes (y-axis: 0.1 and 0.9 are bad and good outcomes, respectively) in each trial. For visualisation purposes, the two 30 trials long blocks of each of the four block-types were concatenated. While related stimuli in +Corr blocks (right panels) are associated with exactly the same probability, their corresponding light and dark grey lines are slightly offset for visualisation purposes.

### Behavioural Modelling

We modelled the subjects’ behaviour using an adapted delta-rule, with a crucial addition of “cross-terms” that enable learning from one stimulus to another. Note this is not intended to be a process model of how the brain solves the task. It is intended as a descriptive model that allows us to test whether, and to what extent, subjects are guided by the relational structure of the task. The model separately tracks the probabilities of a “good” outcome associated with the three stimuli, all of which are updated every trial. Following an outcome at trial *t* where stimulus *X* was presented, the estimate of the outcome probability associated with *X* is updated according to the classic delta-rule:

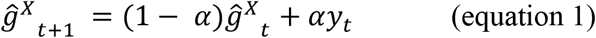

Where 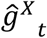 is the outcome probability estimation for stimulus *X* before trial *t, α* is the learning rate, and *y_t_* ∈ {−1,1} is the binary outcome at trial *t*.

Crucially, the estimates of the probabilities associated with the two other stimuli are also updated:

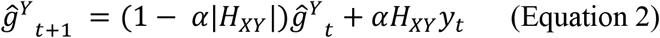

Where *H_XY_* is the cross-term between stimuli *X* and *Y*, fitted to the subject’s behaviour in each block. *H_XY_* = 1 indicates subjects treated *X* and *Y* as the same stimulus for learning purposes - for +Corr stimuli pairs this is the correct correlation structure. Similarly, *H_XY_* = −1 and *H_XY_* = 0 indicates correct correlation knowledge for -Corr and 0Corr pairs, respectively. *y_t_* is still the outcome in trial *t* (a trial where stimulus *X* was presented).

We henceforth refer to the full model, including the 3 cross-terms, as STRUCT. We also used a structure-naïve Rescorla-Wagner model (equivalent to setting the cross-terms in the STRUCT model to 0), which we refer to as NAÏVE.

### Subjects’ behaviour

To determine whether subjects used the relational structure to inform their decisions we employed two approaches: a cross-validation approach and a cross-terms analysis approach. In the cross-validation analysis, for each subject we separately fitted the parameters of the STRUCT and NAÏVE models to data from the 4 block-types, treating trials from each block-type as a separate training set. This resulted in 8 fitted models: 4 STRUCT models and 4 NAÏVE models. We then performed cross-validation by testing how well each of the 8 models predicts subjects’ choices in all 4 datasets (Figure 2a). As expected, STRUCT models generalised much better when trained and tested on different datasets of the same relational structure (despite different stimuli, squares highlighted in pink), than when trained on one relational structure and tested on the other (squares highlighted in green; comparison of sum(-log(likelihood)) of pink vs green elements; two-tailed paired t-test, t(27) =9.54, P<10^-9). Notably, the within-structure cross-validated STRUCT models (squares highlighted in pink) performed better than the NAÏVE models, even when the latter were trained and tested on the same data (squares highlighted in grey; compare pink vs grey elements in Figure 2a: two-tailed paired t-test, t(27) =4.29, P=0.0002). To further demonstrate this effect, we plotted the histograms of outcome probability estimations 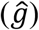 from the relevant within-structure cross-validated STRUCT model (pink elements in Figure 2a; Figure 2b, left) and the NAÏVE model trained and tested on the same data (grey elements in Figure 2a; Figure 2b, right), split by subjects’ choices. When the models made different predictions, subjects tended to choose in accordance with the within-structure cross-validated STRUCT models (Figure 2b). These results suggest that sensitivity to the correlation structure, here afforded by the cross-terms of STRUCT models, is needed to capture subjects’ behaviour.

Due to the small number of trials in each block-type, for the analyses in the rest of the manuscript we trained STRUCT and NAÏVE on each subject’s concatenated data from all blocks of the same structure. This resulted in 10 (5 parameters x 2 structures) and 4 (2 parameters x 2 structures) fitted parameters per subject for the STRUCT and NAÏVE models, respectively. The fitted cross-terms of the STRUCT model indicated that subjects indeed used the correlation structure correctly (Figure 2c; Corr vs 0Corr (mean across all blocks of |*H_AB_*^−,+^| vs (|*H_AC_*| + |*H_BC_*|)/2), one-tailed paired t-test, t(27) = 13.06, P<10^-12). The STRUCT model explained subjects’ behaviour better than the NAÏVE model, even when accounting for the extra free parameters (see a formal model comparison in Table S2). Subjects benefited from the extra information afforded by the correlation structure, and performed better in trials of one of the two related stimuli than in the control stimulus trials (Figure S2a). There were no significant differences in reaction times between trials under the three possible correlation types (Figure S2b), and no effect of task switching costs on reaction times between related and unrelated pairs of stimuli (Figure S2c). These results indicate that the subjects who were scanned indeed used the relational structure.

**Figure 2.**
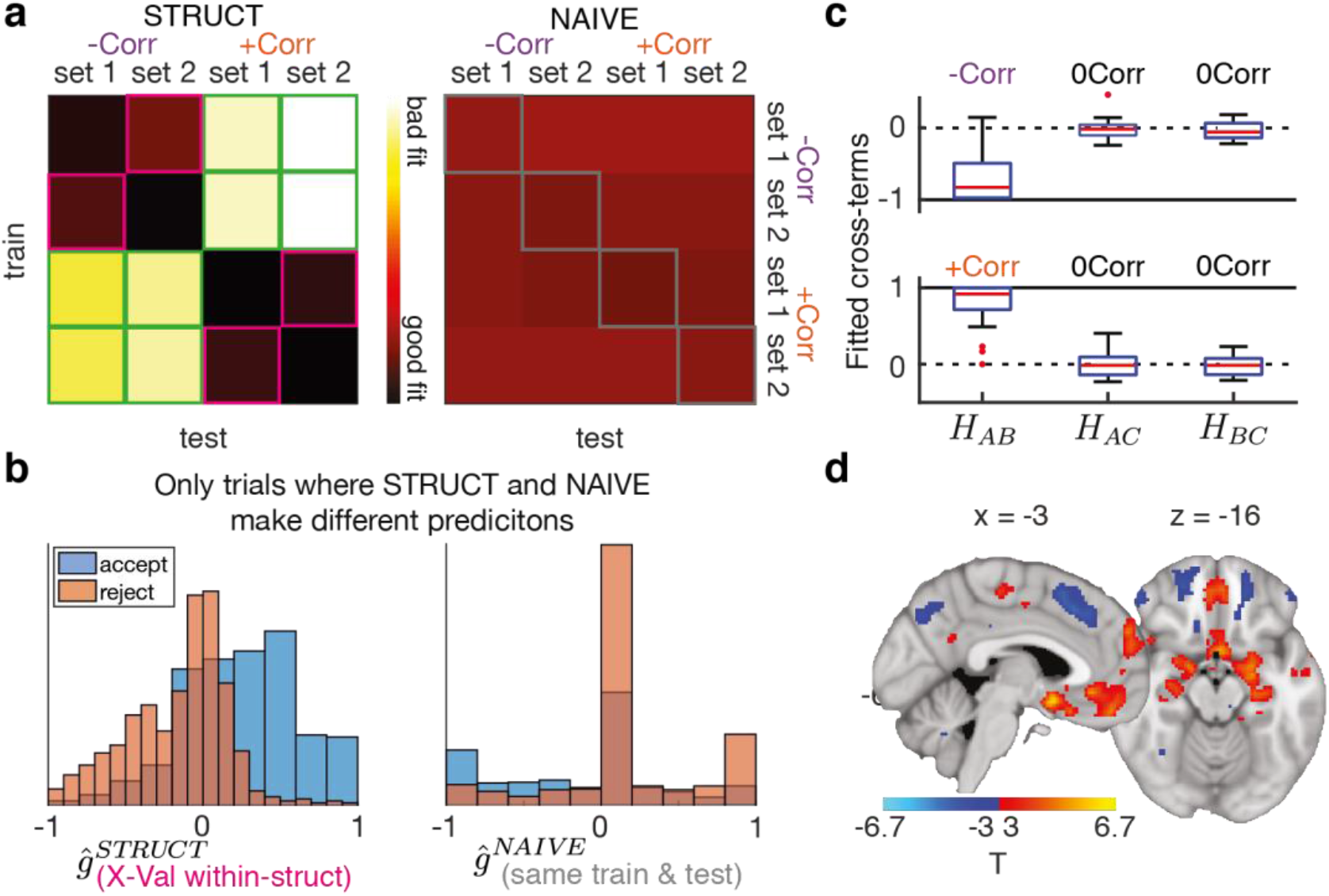
Subjects use the correlation structure correctly. **a.** Negative log likelihoods for STRUCT (left) and NAÏVE (right) models (same scale for both matrices). Pink elements: STRUCT models, cross-validated within-structure. Green elements: STRUCT models, cross-validated across structures. Grey elements: NAÏVE models, trained and tested on the same data. **b.** Histograms of the estimated outcome probabilities for trials where subjects accepted (blue) or rejected (orange). Left: STRUCT models trained on data with the same structure but different stimuli set (pink elements in **a.**). Right: NAÏVE models, trained and tested on the same data (grey elements in **a.**). Histograms only include trials where the models make different predictions. **c.** Fitted cross-terms for pairs of stimuli in all -Corr (top) and +Corr (bottom) blocks. Red central line is the median, the box edges are the 25^th^ and 75^th^ percentiles, the whiskers extend to the most extreme datapoints that are not considered outliers, and the outliers are plotted as red circles. **d.** Effect of the chosen action value estimates from STRUCT model, in a GLM where it competes with estimates from NAÏVE model (replication of Hampton et al^23^).

### The reward network and the hippocampus use the relational structure to encode the value of the chosen action

Our first aim for the analysis of the fMRI data was to replicate a previous report by Hampton et al.^23^, who found evidence of knowledge about the relational structure of the task in known neural signals of RL. We compared how well a model that uses relational structure (STRUCT) explained neural signals relative to one that does not use structure (NAÏVE). In both models, we calculated the value of the chosen action (accept/reject) on each trial of the two related stimuli (A and B). Note that the chosen action value has the same magnitude as the stimulus value tracked by the models but has an opposite sign on rejected trials. The chosen action value estimates were used to construct one regressor per model at the time of stimulus presentation (GLM1). Estimates from both models were pitted against each other in the same GLM, meaning any variance explained by a particular regressor was unique to that regressor, allowing us to compare the neural signals uniquely explained by each model. The contrasts in this section only included the two related stimuli in each block, as the differences between the models for the control stimulus were negligible.

A network of regions including the medial prefrontal cortex (mPFC), the amygdala (AMG), the anterior hippocampus (HPC) and the EC coded positively for the chosen action value from the STRUCT model, while a network including the anterior cingulate cortex (ACC), insula, angular gyrus and most of the orbitofrontal cortex (OFC) showed negative coding (Figure 2d & Table S3; Similar results were obtained for a GLM that included reaction time as a covariate (Figure S3a) and for the STRUCT > NAÏVE contrast (Figure S3b). These results largely form a replication of the work of Hampton et al^23^. This indicates the reward network and HPC use the relational structure to calculate the value of chosen actions.

### Entorhinal representations generalise across tasks with the same structure but not those with different structures

A representation of the relational structure of the task should be similar (generalise) for stimuli which are part of the same relational structure, but dissimilar for stimuli under a different relational structure. We asked whether any region on the cortical surface displayed these properties at the times of stimulus presentation, using Representational Similarity Analysis (RSA^26^) with a searchlight approach.

A searchlight centred on a cortical voxel consisted of the 100 surrounding voxels with the smallest surface-wise geodesic distance from the central voxel. For each searchlight, we obtained 16 patterns of (whitened within-searchlight) regression coefficients of the responses to presentations of each of the two related stimuli (A and B) in each of the 8 blocks (GLM2). In other words, we obtained two patterns, one from each of the repeated runs, for each of our 8 conditions (a particular A or B stimulus under a particular correlation structure). To control for effects of time, we used a “cross-run correlation distance” where only patterns from different runs (i.e. more than 30 minutes apart) were correlated with each other. That is, to define the distance *d_i,j_* between conditions *i* and *j* we first calculated the correlation distance (1 – *r*) between the condition *i* pattern from run 1 and condition *j* pattern from run 2, and then calculated the correlation distance between the condition *j* pattern from run 1 and condition *i* pattern from run 2. *d_i,j_* was defined as the mean of these two distances. Notably, this means that the diagonal in the symmetric Representational Dissimilarity Matrix (RDM) is meaningful, and shows the consistency between the two runs of the same condition.

This resulted in an 8 conditions by 8 conditions symmetric RDM, summarising the representational geometry in the searchlight (e.g. Figure 3b). The ideal structural representation can be formalised as an 8×8 model RDM, where the distances between conditions are determined by relational structure (Figure 3a). To test whether the data RDM of a given searchlight was consistent with the model RDM, we calculated the contrast between the means of the data RDM’s hypothesised “dissimilar” and “similar” elements (white and black elements in Figure 3a, respectively). We verified that this contrast did not correlate with possible behavioural confounds such as reaction time, correctness or task witching costs (Figure S2d, left). We then used permutation tests to ask whether this contrast was significantly positive across subjects (see Methods). We repeated this procedure for each searchlight centre on the cortical surface, resulting in a cortical map of p-values.

The only cluster to survive multiple comparisons correction across a cortical hemisphere was located focally in the right entorhinal cortex (Figure 3b, 3c and 3d, top, P=0.045 FWE corrected for cluster mass, cluster-forming threshold P<0.001, peak MNI coordinates: [25,-5,28]). This effect did not change when we repeated the analysis using model RDMs where same-stimuli or same stimuli set elements were ignored (Figure S4a and S4b). The effect was therefore not driven by background colour or low-level plasticity between stimuli that appear in the same block, but rather by a representation of the relational structure between the stimuli in the task. We note that a cluster in the right temporal-parietal junction (TPJ), where evidence of grid-like coding has been reported 18,19, was the strongest and largest amongst the few other clusters surviving the cluster-forming threshold of P<0.001 (but did not survive FWE-correction). To verify that our analysis approach was indeed valid, we tested for stimulus visual identity coding using a similar procedure with an appropriate model RDM (Figure 3a, bottom). As expected, we observed bilateral effects in visual areas, peaking in the lateral occipital cortex (Figure 3b, 3c and 3d, bottom, P<0.05 FWE corrected on cluster level, cluster-forming threshold P<0.001, peak MNI coordinates: [44,-74,-4]). Therefore, the relational structure of our task is represented and generalised in EC.

**Figure 3.**
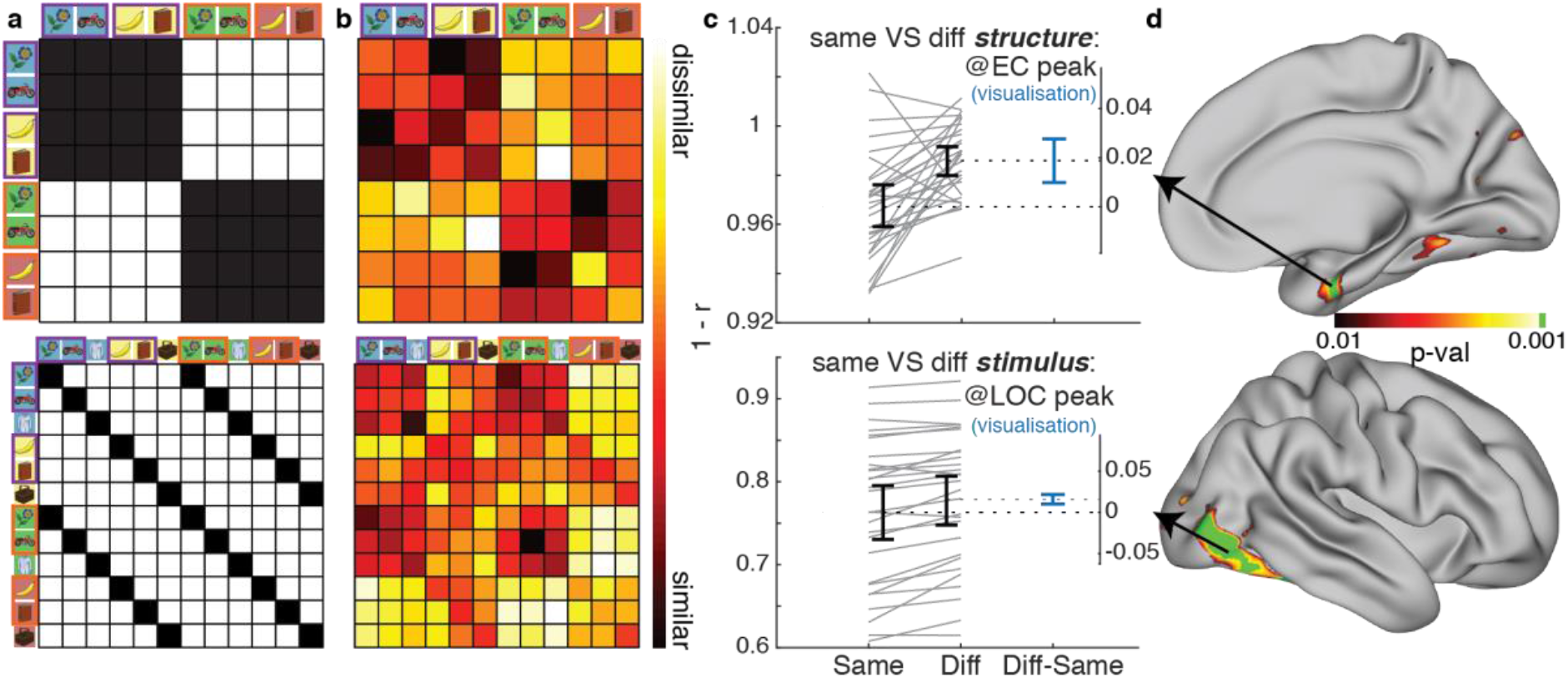
The relational structure of the task is represented in the entorhinal cortex. Top: relational structure effect, peaking in EC. Bottom: stimulus identity effect, peaking in LOC. **a.** Model RDMs. Black elements should be similar, white elements should be dissimilar. Pairs of stimuli with purple and orange rectangles around them are -Corr and +Corr, respectively. **b.** Visualisation of the data RDM from peak vertex of the effect, marked with an arrow in **d**. **c.** Visualisation of the paired mean difference effects between *same* (black RDM elements in **a.**) and *different* (white elements in **a.**) pairs of conditions, from the peak vertex of the effects. Both groups are plotted on the left axes as a slopegraph: each paired set of observations for one subject is connected by a line. The paired mean difference is plotted on a floating axis on the right, aligned to the mean of the *same* group. The mean difference is depicted by a dashed line (consequently aligned to the mean of the *diff* group). Error bars indicate the 95% confidence interval obtained by a bootstrap procedure. **d.** Whole surface results, right hemisphere. Clusters surviving FWE correction across the whole surface at a cluster forming threshold of P<0.001 are indicated in green.

### vmPFC and ventral striatum represent the relational structure in learning signals

The vmPFC is central to reward-guided learning^22^. Intriguingly, previous studies suggest grid-like coding can also be found in mPFC^18–21^. We hypothesised that vmPFC learning signals of the type typically observed in RL tasks^23–25^ may be sensitive to the relational structure. That is, because prediction errors have different implications for learning under the two different relational structures, we hypothesized that vmPFC would not simply encode a signal that monotonically increases with prediction error as previously reported^23–25^. Instead, we hypothesized that the representation of prediction errors across voxels would differ depending on the relational structure.

We first found a strong univariate prediction error signal in a network of regions including vmPFC (Figure 4b, inset) and the ventral striatum (vStr, Figure 4c, inset), in line with previous findings^23–25^. Note that the prediction error used here is the “correctness” prediction error, defined as the magnitude of the prediction error from the STRUCT model, and a sign that depends on the congruence between the subject’s choice and the outcome: positive when the outcome matches the subject’s choice (accept-“good” outcome; reject-“bad” outcome), and negative when choice and outcome are incongruent (accept-“bad” outcome; reject-“good” outcome).

We next asked whether the multivoxel pattern of this prediction error signal depends on the relational structure on a fine-grained scale, in a multivariate analysis. We conducted a searchlight RSA analysis similar to the one from the previous section, with two notable differences: A) the patterns used as inputs to the RSA were not the average responses to the stimuli, but instead the regression coefficients of the prediction errors on the two related stimuli (A and B) in each block. This means the patterns entering this analysis are the local spatial variations in the representation of prediction errors. B) These analyses only have a single measure per block (prediction error coefficients) as opposed to two (separate stimuli) in the previous section (GLM3). The resulting RDMs are therefore 4×4, not 8×8 (Figure S5a-b).

Because we are testing the multivariate differences on the (orthogonal) univariate prediction error effect, we could use the peaks of the univariate effect to constrain our regions of interest. The top three univariate peaks were in bilateral vmPFC (left hemisphere (LH) peak MNI [-4,44,-20], t(27)=9.77, inset of Figure 4b and Figure S5c left; right hemisphere (RH) peak MNI [8,44,-11], t(27)=10.4, inset of Figure 4c and Figure S5c right) and ventral striatum (vStr, peak MNI [-10,8,-12], t(27)=11.28, inset of Figure 4d). In searchlights centred on these peaks, the multivariate prediction error x structure interaction effect was significant (LH vmPFC: P=0.014, Figure 4a, 4b, Figures S5b and S5c left; RH vmPFC: P=0.02, Figures 4c and S5c right; vStr: P=0.034, Figure 4d). We also note that other regions where grid-like coding was previously observed, such as the posterior cingulate cortex (PCC)^18,19^ showed this interaction effect, as well as other reward-related regions like the ventrolateral PFC^22^. See Figure S5c for a description of these exploratory results. As with the relational structure effect, we verified its interaction with prediction errors did not correlate with any behavioural confound (Figure S2d, middle).

These results indicate that prediction error signals in vmPFC and vStr (and perhaps vlPFC and PCC) depend on the current relational structure of the task. The critical difference between the two relational structures in our experiment is not how the prediction error should be computed, but rather how it should be used to inform future behaviour – how should ‘credit’ for the error be assigned^1,3,27–30^. One intriguing possibility is therefore that different representations of prediction errors allow different credit assignment for the same prediction errors in the two relational structures. Factorising these computations from the sensory particularities of the task allows them to be rapidly ‘remapped’ to new stimuli for rapid learning.

**Figure 4.**
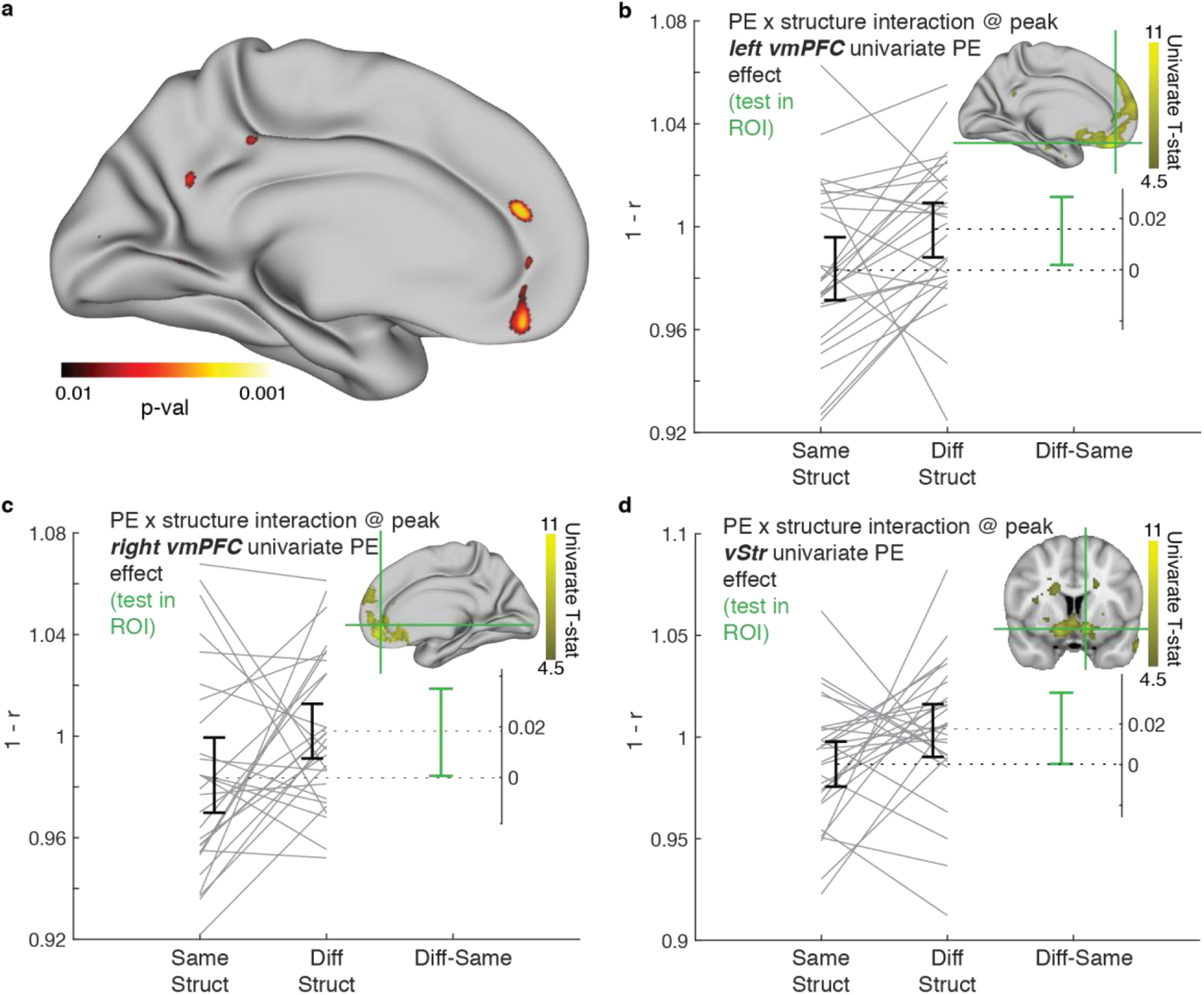
Prediction error signals in vmPFC and ventral striatum depend on the current relational structure of the task. **a.** Visualisation of whole-surface results of the multivariate prediction error x relational structure interaction effect, medial left hemisphere. **b.** Interaction effect at the left hemisphere vmPFC peak of the univariate prediction error effect (MNI: [-4,44,-20]). **c.** Interaction effect at the right hemisphere vmPFC peak of the univariate prediction error effect (MNI: [8,44,-11]). **d.** Interaction effect at the ventral striatum peak univariate prediction error effect (MNI: [-10,8,-12]). Brain images in the insets of **b, c and d** show the univariate prediction error effect (projected on the surface in **b and c**). Legend for panels **b, c** and **d** is the same as in **Figure 3c**.

### OFC representation of task space respects the relational structure

The precise computations that mediate model-contingent credit assignment depend on the brain’s internal representation of the task state-space^1–3,30,31^. In many laboratory tasks, including ours, more than one task representation is possible^1,31^. However, to benefit from the relational knowledge, any such representation must differentiate the related stimuli (A,B) from the control stimulus (C) so that AB outcomes can cause learning on both A and B but not C. For example, one possible representation is to maintain a single variable encoding the state of the AB pair^1^ and a second variable encoding the state of C. Such a representation would be compatible with a representation of the current task state, which has recently been suggested to reside in the OFC^1,2^.

We therefore tested for regions where the representations of AB trials were more similar than AC or BC, again by constructing RDMs. To increase statistical power, we consider representational similarity at stimulus time (S-S), at outcome time (O-O) and between stimulus and outcome (S-O). It is particularly interesting for credit assignment if representations generalise from outcome time to the time when the stimulus was presented. Note that unlike previous analyses, these RDMs are defined within blocks, not between blocks, made possible as ABC trials are presented in pseudo-random order. Hence, within each block, we used correlation distances to calculate three 3×3 data RDMs (S-S, O-O, S-O), where the three conditions in each RDM correspond to the three stimuli. We then averaged the RDMs across all 8 blocks, and defined contrasts between unrelated (AC, BC) and related (AB) pairs of stimuli. Finally, we averaged the three contrasts to obtain a single measure of the latent AB variable representation, and submitted it to permutation tests as described above.

Based on previous suggestions^1,2^, we focused on the OFC as an ROI for this effect. Indeed, the strongest vertex-wise effect in cortex (Figure 5a) was observed in the left OFC (peak at MNI [-18,30,-22]), and OFC clusters survived small volume correction bilaterally (FWE-corrected for cluster mass with a cluster-forming threshold of P<0.01 in an OFC mask: left Brodmann areas (BAs) 11,13,14: P = 0.013; right BA 11 P=0.04). All three experimental periods RDMs (S-S, O-O, S-O) contributed to the OFC effect (Figure 5b and Figure S6b-d). We also observed an unexpected effect in the right posterior parahippocampal (pPHC) gyrus (P=0.034 whole-surface FWE-corrected for cluster mass with a cluster-forming threshold of P<0.01). See Methods for description and discussion of statistical correction.

**Figure 5.**
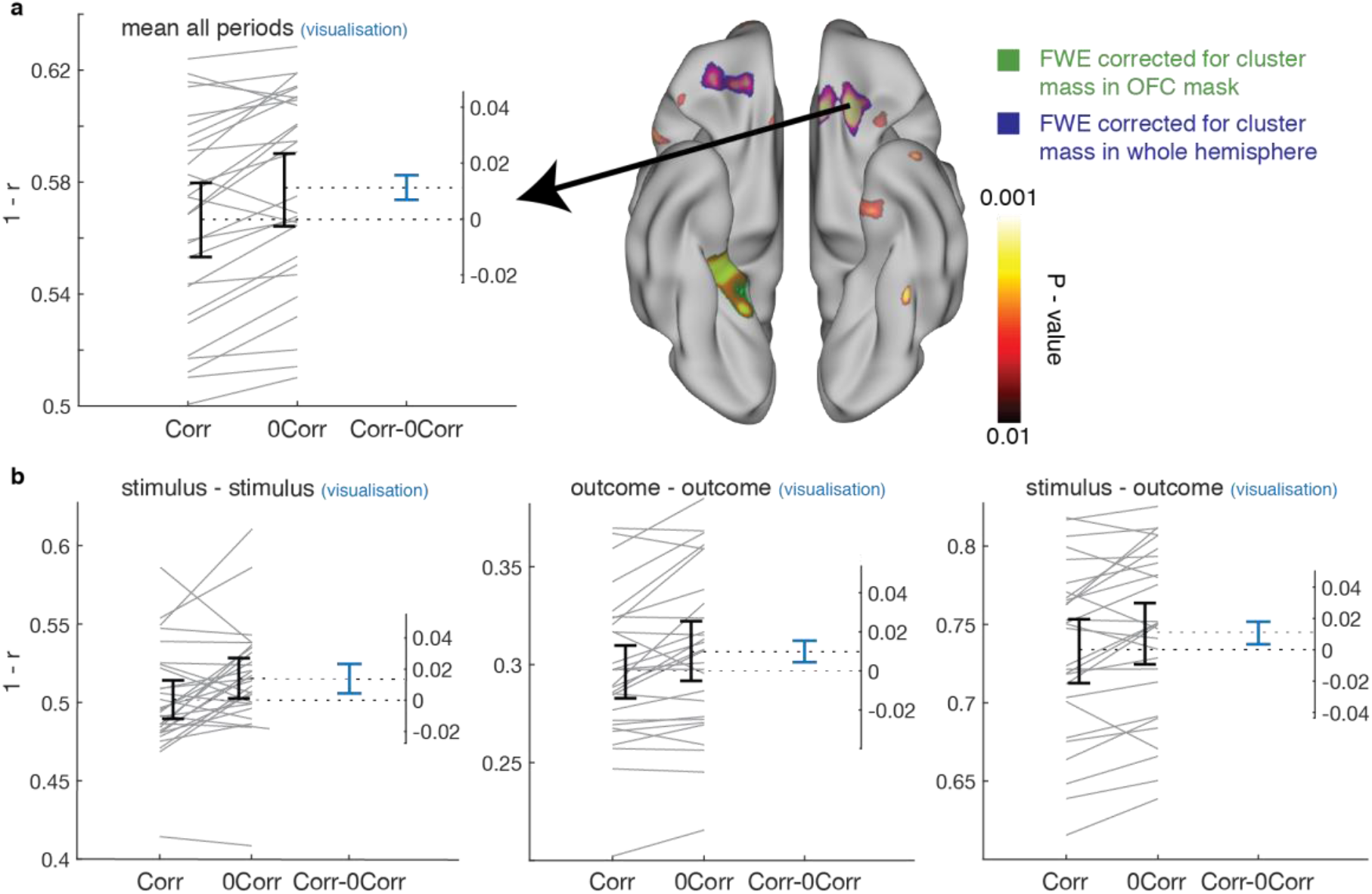
OFC represents a latent variable, respecting the task structure. **a.** Right: Whole-surface results (orbital view) for the [unrelated stimuli – related stimuli] contrast, averaged over all experimental periods (S-S, O-O, S–O). The semi-transparent uncorrected P-value map is overlaid over the clusters that survived FWE correction: the OFC clusters in blue survived small volume correction in an OFC mask, while the pPHCg cluster in green survived correction in the entire hemisphere (see discussion of statistical correction in the Methods). Left: Visualisation of the effect from the peak of the OFC effect, located in the left OFC (MNI [-18,30,-22]). **b.** Same as **a. (left),** separated to the different experimental periods (S–S, O-O, S-O). Legends are the same as in **Figure 3c**.

## Discussion

Understanding how the brain represents abstract task state-spaces remains a major challenge. Our data joins growing evidence suggesting state-space representations rely on the hippocampal formation^4,8,32–34^ and interconnected regions in the ventral PFC^1,2,34,35^. This is particularly interesting in light of the historical role of these regions in generalisation and relational reasoning^36–41^, which are essential for efficient task representations. Here, we show that these regions, and in particular the same areas where evidence of grid cells was found, generalise the relational structure of a non-Euclidian RL task. EC representations of stimuli generalised over tasks with the same structure, but not over tasks with a different structure. The same was true for vmPFC (and vStr) representations of prediction error. These results suggest a common framework for the representation and generalisation of task structures in a wide variety of domains. In addition, we found that OFC and posterior PHC gyrus representations were consistent with the coding of a behaviourally-relevant latent variable, respecting the task structure^1,2^.

Our experiment can be viewed as a set of non-spatial “remapping” experiments. The vertical arms in Figure 1b (same task structure, different stimuli) are analogous to the classic spatial sensory remapping experiments in rodents^12,14^, where an animal is moved between different sensory examples of the same task structure (namely free foraging in a Euclidean 2D space). In these experiments, entorhinal grid cells maintain (generalise) their covariance structure across environments (i.e. they do not remap, in contrast to hippocampal place cells^42^). Such generalisation was also observed in Macaque OFC neurons, across different sensory examples of an economic decision-making task^43^. The horizontal arms in Figure 1b (same stimuli, different task structure) can be viewed as “task remapping” experiments. In the spatial case of such experiments, where animals are required to perform different tasks in the same sensory environment, recent evidence suggests the grid code changes (remaps) across tasks^44,45^. Our EC effect (Figure 3) mirrors these results in a non-spatial RL task in humans. Future work might address whether the exact same neuronal population underlies generalisation in both spatial and non-spatial tasks.

We believe the comparison to grid cells is essential to understand these results: First, grid cells abstract and generalise the relational structure of 2D environments. This is demonstrated in remapping tasks^7,12,14^, which inspired the design of our task. Second, an important property of the relational structure that they generalise is the correlation structure of observations, that is shared between all 2D environments, even when the observations themselves differ. Here, we similarly manipulate the correlation structure of observations, only these observations are rewards in an abstract RL task rather than locations in a spatial task. Third, grid cells reside predominantly in the entorhinal cortex, but also in human mPFC (and other areas) – the areas where we find our effects. However (and crucially), we do not claim that hexagonal grid-like patterns^18,19,21^ underlie our effects, nor that it is possible to measure these here. A hexagonal representation, while being the most efficient representation for generalising the relational properties of 2D Euclidean space^9,10^, would not be useful for generalisation in our task. Rather, we test - in a non-Euclidean RL state-space - for the underlying computational functions (reviewed in Behrens et al.^4,6,7,9^) that lead to the hexagonal pattern in 2D space.

A unified framework for the representation of task structure might also afford a new way to interpret standard RL neural signals like prediction error. While the dependence of prediction error signals on prior, unobservable information has been reported previously^46^, here we report a novel relational aspect to this context dependency. Prediction error representations that were used in different ways (opposite update signs of the related stimulus for -/+Corr blocks) differed anatomically on a fine-grained level. Our vmPFC prediction error x structure interaction effect (Figure 4) brings together observations about vmPFC function from several seemingly disparate fields, including RL^22^. Patients with vmPFC lesions show a selective valuation deficit when value comparison should be based on attribute configuration, i.e. when relationships between object elements are important for the valuation^47^. In the memory literature, vmPFC has been strongly implicated in the representation of schemas – abstract structures of previous knowledge, which bear many parallels to the relational structures discussed here^48,49^. vmPFC is particularly important when new information is assimilated into an existing schema^50,51^, analogous to a prediction error update of the internal model of the task within the current structure. Finally, some of the strongest effects of spatial and nonspatial grid-like coding in fMRI were recorded in the medial PFC^18–20^. Notably PCC, where we found the strongest prediction error x structure interaction effect in our exploratory analysis, also exhibits grid-like coding^18,19^ and has also been strongly implicated in the representations of schemas^49,50^. Both the vmPFC structural learning signal and the latent variable representation in OFC^1–3,30^ might be useful for facilitating appropriate model-contingent credit assignment^27–29^.

Though our manuscript focuses on novel data regarding structural representations of task elements (Figure 3) and learning signals (Figure 4), we also report a key replication (Figure 2d). This is notable, as the effect that is replicated is subtle - the unique contribution of model-based (over model-free) value in a relational RL task – and yet the effect replicated in detail. Although the original paper by Hampton et al.^23^ focuses on the vmPFC, the whole-brain results for the relevant contrast can be seen in their Figure 2a. We also report the negative effects of this contrast (in angular gyrus, ACC and OFC, where negative correlates of value are often observed). These negative effects do not form part of the replication, as Hampton et al. did not report negative effects (though it is plausible they existed in their data).

Learning can be dramatically improved by a useful representation of the world you are learning about. Here, we show that the brain can “recycle” (generalise) these representations, enabling fast and flexible inferences. We believe the comparison of generalisable representations in our abstract RL task to their parallels in spatial cognition is a useful one, and can suggest a path for a more precise understanding of the nature of these representations.

## Acknowledgments

We thank Jacob Bakermans, Anna Shpektor, Philipp Schwartenbeck, Avital Hahamy and Shirley Mark for useful comments on earlier versions of the manuscript.

## Author contributions

A.B.B, T.H.M and T.E.J.B designed the study. A.B.B acquired the data. A.B.B, T.H.M, M.G, H. N and T.E.J.B analysed data. A.B.B and T.E.J.B wrote the manuscript.

## Methods

All code is a available at https://github.com/alonbaram2/realtionalStructure_NN2020. Data is available at ****. Unthresholded statistical maps can be obtained at https://identifiers.org/neurovault.collection:7150

### Subjects

We trained 49 volunteers over 4 days on an online version of the task. 17 subjects did not proceed to be scanned as they either did not comply with task demands (e.g. failed to complete training on time) or did not reach a behavioural criterion for knowledge of the outcome probabilities correlation structure (a difference of more than 0.3 between the fitted cross-term of the related stimuli and the mean of the fitted cross-terms of the unrelated stimuli, see below). 32 volunteers (aged 21-32 years, mean age 23.4, 18 females) with normal or corrected-to-normal vision and no history of neurological or psychiatric disorders participated in the fMRI experiment. 4 subjects were excluded from the analyses: 3 due to technical difficulties during the scanning, and one due to excessive motion. Hence, all analyses presented are based on data from 28 subjects. All subjects gave written informed consent and the study was approved by the University of Oxford ethics committee (reference: R51215/RE001).

### Training and task

Subjects trained online for 4 days prior to the scan day, and were scanned on day 5. In each training session, subjects performed a task where three 1-armed bandits were interleaved pseudo-randomly. The bandits were cued by three different visual stimuli, randomly sampled without replacement for each session from a bank of 35 images. Two of the bandits (bandits A & B) had correlated outcome probabilities, while the third (C) was independent. There were two possible correlation structures for the outcome probabilities of bandits A & B: positive correlation (+Corr blocks) or negative correlation (-Corr blocks). Each online training session comprised of 8 blocks with 60 trials each (3 stimuli X 20 trials per stimulus).

In each trial, subjects viewed one of the three stimuli and had to indicate their prediction for its associated binary outcome (a “good” or a “bad” outcome, demarked by a King of Hearts or Two of Spades card, respectively) by either accepting or rejecting the stimulus. Thus, there was always one correct answer in each trial: subjects should accept a stimulus if they predict the outcome to be the “good” outcome, and should reject if they predict the outcome to be the “bad” outcome (Figure 1a). Only in accepted trials, the subject would either win or lose a point, depending on the outcome (i.e. accepting incorrectly resulted in a loss of a point). Outcome identity was revealed in all trials, including rejection trails (except on training days 3 and 4, see below), even though the subject’s score did not change in these trials (Figure 1a). Predictions of the outcomes could be formed based on the recent history of outcomes. The outcome probabilities switched pseudo-randomly between 0.9 and 0.1 with an average switch probability of 0.05 in each trial of the training sessions. As the two correlated bandits switched together, subjects could use their knowledge of the correlation structure to learn from the outcome on stimulus A about stimulus B and vice versa (Figure 1c). Subjects were informed and reminded at the beginning of each training block that two of the bandits had correlated outcome probabilities, and that this correlation might be positive or negative. However, subjects had to infer which two bandits were correlated and what the sign of the correlation was.

The training schedule for an example subject is shown in **Table S1**. On day 1 subjects completed two sessions with a different triplet of stimuli used as the ABC cues in each session. In both sessions, two of the stimuli had a particular correlation structure, counterbalanced across subjects. That is, half of the subjects performed two sessions of 8 - Corr blocks each, while the other half performed two sessions of 8 +Corr blocks. Day 2 was identical to day 1, except that the two stimuli sets used were novel and the correlation structure was the one that the subject did not experienced on day 1. On day 3 subjects again completed 2 sessions with a novel stimuli set per session, where the correlation structure between two of the stimuli alternated between blocks. The correlation structure was indexed by the background colour of stimuli (e.g. Figure 1b and 1c), and subjects were informed that the combination of stimuli set, background colour and correlation structure in day 3 will be the same in days 4 and 5, including in the scanning task. Thus, subjects could already learn the background colour-correlation structure mapping prior to entering the scanner. On day 4 subjects completed one session with all the 4 possible block-types (2 stimuli sets x 2 correlation structures, e.g. Figure 1b and 1c). To reduce available information and facilitate subjects’ need to use the extra information afforded by the correlation structure, no counterfactual feedback was given on rejection trials in any of the last 15 trials of a block in training days 3 and 4.

Prior to scanning on day 5, subjects completed a pre-scanning reminder session with all 4 block-types, again with the same stimuli set - background colour - correlation structure combinations as in days 3 and 4. In both the pre-scanning session and during scanning, full outcome feedback was given, including in all rejection trials. During scanning, subjects completed 8 blocks of 30 trials each (10 trials per stimulus), with a break after 4 blocks for structural and field-maps scans. The two groups (runs) of 4 blocks included one of each of the 4 experimental block-types in a pseudo-random order, counterbalanced across subjects. Outcome probabilities switched faster in the scanner than during training due to the shorter blocks, with a switch probability of 0.15.

Before the first trial of each block, all three stimuli and the background colour of that block were presented. A trial was on average 11.5 seconds long, progressing in the following order (Figure S1): 1) A stimulus would appear in the middle of the screen, together with the available choices, e.g. left for accept (corresponding to an index finger button press) and right to reject (middle finger button press). The left/right mapping to choices was counterbalanced across subjects but stable within-subject. 2) After 1.5 seconds, the frame of the stimulus and the accept/reject text turned white, indicating that choice can now be made. Subjects who indicated their choice prior to the appearance of the white frame lost half a point. 3) Subjects indicated their choice by pressing either the index or middle finger buttons. Next, a red rectangle appeared around the chosen option for 0.5 seconds. 4) A white fixation cross appeared for a variable period, drawn from an exponential distribution with a mean of 4.5 seconds and truncated between 3.5-5.5 seconds. The purpose of this long delay between choice and outcome was to enable the independent analysis of both periods, due to the sluggish nature of the hemodynamic response function. 5) The outcome of the trial appeared for 1 second in the middle of the screen - either the “good” outcome (King of Hearts card) or the “bad” outcome (Two of Spades card). If the subject has accepted the trial earlier, they will either win or lose a point: if they accepted correctly (outcome was “good”), a “sack of gold” image would appear in the left side of the screen to indicate that a point was gained; if they accepted incorrectly (outcome was “bad”), a “no sack of gold” image would appear to indicate that a point was lost. If the subject rejected the trial, no points would be won or lost (and hence no “sack of gold” or “no sack of gold” image would appear), but the outcome card image would still appear (Figure 1a). Hence subjects received full counterfactual feedback. Note that subjects should reject trials if they predict the outcome to be the “bad” outcome, as they will lose a point for incorrectly accepting a trial. 6) A white fixation cross appeared for a variable inter trial interval, drawn from an exponential distribution with a mean of 3 seconds and truncated between 2.5-4 seconds.

### Behaviour modelling

We modelled the behaviour of the subjects using an adapted delta-rule model [1]. The original delta-rule model estimates the outcome probabilities for a given stimulus using the following equation:

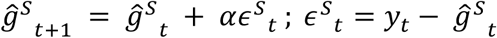, where *S* ∈ {*A, B, C*}; {*A, B, C*} are the three stimuli presented in a block, 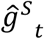 is the value estimation of stimulus *S* before trial *t* (which can be thought of as the “good” outcome probability estimation for stimulus *S* before trial *t*, mapped to the [-1,1] interval instead of [0,1]), *α* is the learning rate, *ϵ* is the outcome prediction error, and *y_t_* is the outcome at trial *t*: *y_t_* = 1 for the “good” outcome and *y_t_* = −1 for the “bad” outcome. Note that the model estimates the stimulus value, based on the outcome identity (which card was obtained - not the reward actually obtained by the subject), and is agnostic to the choice the subject made. The stimulus value should be distinguished from the “chosen action value” used in GLM1 (see below), which has the same magnitude as the stimulus value, but has an opposite sign on rejection trials: in a trial where the subject’s hypothetical estimate of the stimulus value is very low (close to −1), they will be confident in rejecting the trial, making the value of the chosen “reject” action high (close to +1). Similarly, *ϵ* should not to be confused with the “correctness” prediction error used in GLM3 and Figure 4, which has the same magnitude as *ϵ* but with a sign determined by the congruence between the subject’s choice and the outcome. The stimulus value estimation can then be used by a “selector model” to make a choice in the next trial, by using a simple sigmoidal function: *P*(*choice on stimulus S on trial t* + 1 = *accept*) = 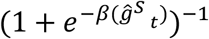, where *β* is the inverse temperature, controlling the randomness of the choice. In this model, *α* and *β* are free parameters fit to subjects’ behaviour.

However, this model does not use any knowledge about the correlations between the outcome probabilities of different stimuli. To allow for this, we added three free parameters to the model, which we refer to as cross-terms. These parameters determine how information on one stimulus affects the outcome estimate on another stimulus.

Following an outcome on stimulus *A*, we update the outcome estimates of all three stimuli:

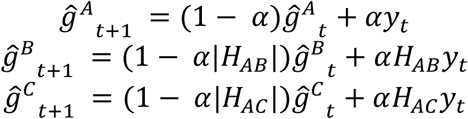

Where −1 ≤ *H_XY_* ≤ 1 is the cross-term for stimuli *X* and *Y*. Note that the first equation (update of the estimate for stimulus *A* following an outcome on stimulus *A*) is identical to the update in the original delta-rule model. *H_XY_* = 1 means stimuli *X* and *Y* are treated as the same: the outcome estimates for both stimuli will be updated in exactly the same way following feedback on one of them. *H_XY_* = −1 means stimuli *X* and *Y* are treated as having opposite (anti-correlated) outcome probabilities. *H_XY_* = 0 means the two outcome probabilities are treated as uncorrelated: the outcome estimates for stimulus *X* will not change following feedback on stimulus *Y*, and vice versa. Analogous updates occur when feedback is given on stimuli *B* or *C*.

To establish the robustness of the STRUCT model in explaining subjects’ choices, we first performed a cross-validation analysis. We fitted both STRUCT (5 free parameters per training data: learning rate, inverse temperature and 3 cross-terms) and NAÏVE (2 free parameters per training data: learning rate and inverse temperature) models to training data from the four block-types (2 structures x 2 stimuli sets) separately, concatenating data from the pre-scanning (1 block of 42 trials per block type) and scanning (2 blocks of 30 trials per block type) sessions. Hence, we fitted 4 STRUCT models and 4 NAÏVE models. We then tested the trained models on data of subjects’ choices from either the same (diagonals in Figure 2a) or different (off-diagonals in Figure 2a) block-type, resulting in 4×4 matrices of the negative log likelihood of the test data given the fitted model (Figure 2a). The outcome estimates for all stimuli were reset to 0 at the beginning of each block. We fit the parameters by maximising the negative log likelihood of the data with respect to the parameters using the Matlab function *fmincon*. The learning rate was constrained to be between 0 and 1, the cross-terms between −1 and 1, and the inverse temperature between 0 and 8.

To further demonstrate the robustness of the STRUCT model, we isolated trials where the NAÏVE model, when trained and tested on the same data (grey elements in Figure 2a, right), predicted different choices than the cross-validated STRUCT model (trained and tested on different data but from the same relational structure, pink elements in Figure 2a, left). We plotted histograms the models’ outcome probability estimation, separately for accept/reject choices of subjects.

For the rest of the paper, we used STRUCT and NAÏVE models fitted on data pooled across all blocks of the same structure (collapsing over stimuli sets). That is, we collapsed data from the subject’s scanning session (30 trials per block x 4 blocks per structure) and the pre-scanning session (42 trials per block x 2 blocks per structure), and fitted the parameters separately for +Corr and -Corr blocks. This resulted in a total of 202 trials for each of the structures (+/-Corr). The final STRUCT and NAIVE models had 10 (5 parameters x 2 structures) and 4 (2 parameters x 2 structures) free parameters per subject, respectively. We confirmed that the STRUCT model provided a better fit to the data even when accounting for the extra degrees of freedom through a formal model comparison (Table S2). Finally, we tested whether subjects indeed used the relational structure correctly by performing a one-tailed paired t-test across subjects between the STRUCT model AB cross-term and the mean of AC and BC cross-terms (Figure 2c).

We note that we do not suggest that this model reflects how the brain solves the task. The model is only used as a way to analyse the behavioural and neural data. Specifically, the model allows us to 1) Establish that the relational structure influenced subjects’ behaviour. 2) Compute a proxy to value and prediction error signals, to use in the fMRI analysis.

### fMRI data acquisition and preprocessing

Data were acquired on a 3T Siemens Prisma scanner, using a 32-channel head coil.

Functional scans were collected using a T2*-weighted echo-planar imaging (EPI) sequence with a multi-band acceleration factor of 3, within-plane acceleration factor (iPAT) of 2, TR = 1.235 s, TE = 20 ms, flip angle = 65 degrees, voxel resolution of 2×2×2 mm and a tilt of 30 degrees relative to axial axis. A field map with dual echo-time images (TE1 = 4.92ms, TE2 = 7.38ms, whole-brain coverage, voxel size 3×3×3 mm) was acquired to correct for geometric distortions. Structural scans were acquired using a T1-weighted MP-RAGE sequence with 1×1×1 mm voxels. Preprocessing was performed using tools from the fMRI Expert Analysis Tool (FEAT), part of FMRIB’s Software Library (FSL [2]). Data for each of the 8 blocks were preprocessed separately. Each block was aligned to the first, presaturated image using the motion-correction tool MCFLIRT [3]. Brain extraction was preformed using automated brain extraction tool BET [4]. All data were high-pass temporally filtered with a cut-off of 100 seconds. Registration of EPI images to high-resolution structural images and to standard (MNI) space was performed using FMRIB’s Linear and Non-Linear Registration Tool (FLIRT and FNIRT [3,5]), respectively. The registration transformations were then used to move each blocks’ EPI data to the native structural space, downsampled to 2×2×2 resolution. No spatial smoothing was performed during preprocessing (see below for different smoothing protocols for univariate and multivariate analyses).

### Univariate analyses

Due to incompatibility of FSL with the MATLAB RSA toolbox [6] used in subsequent analyses, we estimated all first-level GLMs and univariate group-level analyses using SPM12 (Wellcome Trust Centre for Neuroimaging, http://www.fil.ion.ucl.ac.uk/spm).

For univariate analyses, contrasts of parameter estimates were smoothed with a kernel of 5mm FWHM before performing group level statistics.

In the following descriptions, [A, B, C] refer to the three stimuli presented in a particular block, where the outcome probabilities associated with A & B was correlated (either positively or negatively, depending on the block).

To test whether known RL signals in the reward network were consistent with a model that used the relational structure, we constructed a GLM where chosen action value estimates from both STRUCT and NAÏVE models were pitted against each other (without being orthogonolised, Figure 2d and S3). <u>GLM1</u> included the following regressors per each block: two main effect regressors of all related stimuli trials ([AB]): one modelling the times of stimulus presentation and modelling outcome times. Two parametric regressors were locked to stimulus presentation times of [AB]: the value of the chosen option from the STRUCT and the NAÏVE models. In addition, the GLM included several other regressors: C trials at stimulus presentation; C trials at outcome; 2 regressors modelling button presses across all stimuli: one modelling all “accept” trials and one modelling all “reject” trials; 6 motion parameters as nuisance regressors; bias term modelling the mean activity in each block. Figure 1e shows the results of the contrast [STRUCT chosen option value] > [baseline]. Figure S3b shows the results of the contrast [STRUCT chosen option value] > [NAIVE chosen option value]. Figure S3a shows the results of the [STRUCT chosen option value] > [baseline] contrast in a separate GLM where we added [AB trials reaction times] as a parametric regressor locked to the time of stimulus presentation.

See below for a description of the univariate STRUCT model prediction error analysis, the results of which are shown in the insets of Figure 4b and 4c.

### RSA analyses

In this section, we first outline the steps common to all RSA [7] analyses, and then describe details which were specific to each analysis. All RSA analyses were conducted as follows: 1) A searchlight[8] was constructed around each cortical voxel, including the 100 cortical voxels with the smallest surface-wise geodesic distance from the central voxel. This was performed using adapted scripts from the RSA toolbox (Nili et al., 2014 original surface-based searchlight scripts written by Joern Diedrichsen & Naveed Ejaz, code available at https://github.com/rsagroup/rsatoolbox). The searchlight definition depends on cortical reconstruction and alignment performed via Freesurfer’s *recon-all* command [9–15]. 2) For each searchlight, first-level univariate GLMs on unsmoothed data were conducted using the RSA toolbox, based on SPM12. Regression coefficients were then spatially pre-whitened within the searchlight using the RSA toolbox. 3) A distance metric was defined to summarise the representational geometry between conditions. The metrics used were either the cross-run correlation distance or a within-block correlation distance (See below). This resulted in data RDMs of size [number of conditions] x [number of conditions]. 4) A hypothesis about the representational geometry was formalised as a contrast between the mean of RDM elements that should “dissimilar” to each other and the mean of RDM elements that should be “similar” to each other. This contrast was calculated in each searchlight, resulting in a single contrast map per subject. 5) The contrast maps of the different subjects were aligned on the common cortical surface using Freesurfer-based scripts adapted from the RSA toolbox. 6) Group-level statistical significance of the contrast (equivalent to a one-tailed t-test) and family-wise error (FWE) correction were performed using permutation tests [16] using PALM [17]. In this procedure, the contrast values from each subject were randomly multiplied by either 1 or −1, following the null hypothesis that the contrasts are symmetric around 0. The test’s statistic was then defined as the cross-subject average of contrast values. This was repeated 10000 times, creating a null distribution of the means. The true value of this mean was then compared to this null distribution. The resulting (uncorrected) p-value map is displayed in figures 3d, 4a, 5a, S4 and S5c as a heat map, at a threshold of P<0.01. See below for a separate section discussing FWE correction. The paired mean difference across subjects between the two groups of RDM elements at particular vertices of interest is visualised in the Gardner-Altman estimation plots in figures 3c, 4b, 4c, 5a and 5b. The figures were generated using an adaptation of the openly available Matlab package DABEST [18]. We make an important distinction between estimation plots containing data from peaks of FWE-surviving clusters, which are subject to selection bias (and are shown for visualisation purposes only; figures 3c, 5a and 5b), and estimation plots with data from unbiased ROIs, which are not subject to selection bias (figures 4b and 4c. In these ROIs an uncorrected statistical test can be performed.

To search for a representation of the relational structure between stimuli in the task (Figures 3 & S4), we conducted a GLM (<u>GLM2</u>) which included the following regressors per each block: 3 main effect regressors ([A],[B],[C]) modelling the times of stimulus presentation; 3 main effect regressors ([A], [B], [C]) modelling the times of outcome presentation; 2 regressors modelling button presses across all stimuli: one modelling all “accept” trials and one modelling all “reject” trials; 6 motion parameters as nuisance regressors; bias term modelling the mean activity in each block. Only the 2 regressors modelling the presentation of the two related stimuli (A&B) in each block were used in the “relational structure representation” analysis (as stimulus C was not part of a relational structure, Figure 3 top and Figure S4), while all stimulus presentation regressors (A, B & C) were used in the “visual stimulus identity representation” analysis (Figure 3, bottom). Each condition (a particular stimulus under a particular structure) had two patterns (100-long vectors of spatially pre-whitened regression coefficients) – one from each independent run. We defined the cross-run correlation distance between each pair of conditions by averaging the following quantities: [correlation distance between condition *i* pattern from run 1 and condition *j* pattern from run 2] and [correlation distance between condition *i* pattern from run 2 and condition *j* pattern from run 1]. This resulted in [number of conditions] x [number of conditions] data RDM for each searchlight (8×8 for the “relational structure representation” analysis; 12×12 for the “visual stimulus identity representation” analysis). We then defined a hypothesis-driven contrast between RDM elements: In the main “relational structure representation” analysis, “different structure” elements should be more dissimilar to each than “same structure” elements. For the control analyses in Figure S4, we ignored elements of the same visual stimulus (Figure S4a) or the same stimuli set (Figure S4b). In the “visual stimulus identity representation” analysis, “different visual stimulus” elements should be more dissimilar than “same visual stimulus” elements. Using the maximum cluster mass statistic [16] for multiple comparisons correction, we report clusters that survived FWE correction at a cluster-forming threshold of P<0.001 within a cortical hemisphere (green clusters in figures 3 and S4).

To test whether standard prediction error signals depended on the relational structure (prediction error x structure interaction analysis, Figures 4 and S5), we first wanted to replicate the previously described ventral striatum (vStr) and vmPFC univariate prediction error signals [19–22]. We conducted a GLM (<u>GLM3</u>) which included the following regressors per each block: 2 main effect regressors ([AB],[C]) modelling outcome times; one parametric regressor of the “correctness” prediction error from the STRUCT model, locked to the time of [AB] outcome presentation; 2 main effect regressors ([AB],[C]) modelling stimulus presentation times; 2 regressors modelling button presses across all stimuli: one modelling all “accept” trials and one modelling all “reject” trials; 6 motion parameters as nuisance regressors; bias term modelling the mean activity in each block. For each subject, we calculated the contrast [AB STRUCT “correctness” prediction error] > [baseline], smoothed the contrast image using a 5mm FWHM kernel, and obtained group-level results using SPM (Figure 4b and 4c, insets). As expected, the two strongest peaks were in vStr and vmPFC. We then used these peaks as (unbiased) ROIs for the multivariate prediction error X structure interaction analysis. To this end, in addition to the surface-based cortical searchlight procedure described above, we also defined 100 voxels long volumetric searchlights within an anatomical mask of the vStr (Harvard-Oxford Subcortical Structure Atlas). Unlike the surface-based searchlight, the analyses for the volumetric searchlight were all performed in MNI152 space. Only the regressor modelling the [AB] “correctness” prediction error in each block was used in the RSA analyses for Figures 4 & S5, resulting in 8 patterns – one pattern per block. We again used the cross-run correlation distance to collapse patterns of the same conditions (a particular stimuli set under a particular structure) across runs, this time to construct 4×4 data RDMs – one condition per block type. Again, we defined a hypothesis-driven contrast between RDM elements, where “different structure” elements should be more dissimilar to each than “same structure” elements. We report the (uncorrected) p-values obtained by permutation tests in the searchlights centred on the vStr and vmPFC peaks of the univariate prediction error effect (Figure 4). We also present the exploratory, uncorrected results across cortex of this effect (Figure S5c).

To test for a representation of the latent AB variable (Figure 5), we once again used the regression coefficients from <u>GLM2</u>. This time, however, we used the regression coefficients from all three stimuli in each block at both stimulus presentation (S) and outcome presentation (O) times. This was enabled by the long (mean ~4.5 seconds, jittered) S-O delay period. As we were testing for a within-block effect, here we used a different distance metric: a simple correlation distance, calculated within each block. Thus, we obtained three 3×3 data RDMs per block, one for each pair of experimental periods (S-S, O-O, S-O), where the 3 conditions in each RDM correspond to the 3 stimuli presented in the block. Note that each element in the S-O RDM is an average of two correlations, e.g. *d_AB_* = 1 − *mean*(*corr*(*A_o_, B_s_*), *corr*(*A_s_, B_o_*)). We then averaged the RDMs across all blocks (collapsing over blocks of different relational structures). To obtain a single measure of the latent variable representation and to increase statistical power, we averaged the S-S, O-O and S-O RDMs. Next, we defined a hypothesis-driven contrast between RDM elements: unrelated (AC, BC) elements should be more dissimilar to each other than related (AB) elements. We once again used permutation tests for statistical testing and FWE correction (see below for more details).

Lastly, we tested whether the vmPFC prediction error X structure interaction effect correlated on a subject-by-subject level with the mOFC latent variable representation effect, by extracting the contrasts of different subjects at the peaks of both effects and correlating across subjects.

### Multiple comparisons correction

Multiple comparisons correction was performed using the permutation tests machinery (Nichols & Holmes 2002) in PALM [17]: we first thresholded the (uncorrected) P-map at either P<0.001 (Figure 3) or P<0.01 (Figure 5), and measured the mass of all surviving clusters. We then repeated this procedure for each of the 10000 random sign-flip iterations described above, and created a null distribution of cluster masses by saving only the mass of the largest cluster in each iteration. Comparing the true cluster masses to the resulting null distributions results in FWE-corrected P-values.

Whilst the effects in the latent variable representation analysis (Figure 5) do not survive multiple comparisons correction with the same parameters as the entorhinal effect in Figure 3, they do survive with a lower cluster forming threshold of P<0.01 (pPHC gyrus cluster: whole-surface corrected; OFC clusters: small volume corrected in an anatomical mask of OFC derived from the PALS B12 atlas in Freesurfer), indicating that the effects are weaker but larger than the EC effect in Figure 3. This is perhaps not surprising as the OFC and pPHC are larger than EC and therefore can support larger more diffuse clusters. The choice of the OFC as an ROI for correction was based on the prior hypothesis for this effect in OFC, following [23,24]. In the absence of pre-registration, we note that the OFC FWE results in Figure 5, whilst strong, involved minor parameter search and therefore may not control precisely for family-wise errors.

## Supplementary materials

**Figure S1.**
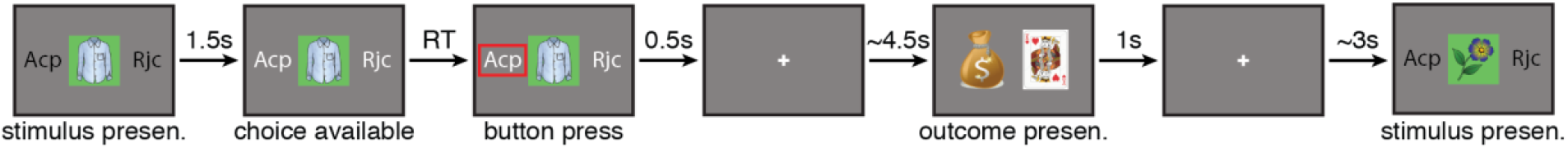
Time course of an example trial. Acp: Accept; Rjc: reject. RT: reaction time.

**Figure S2.**
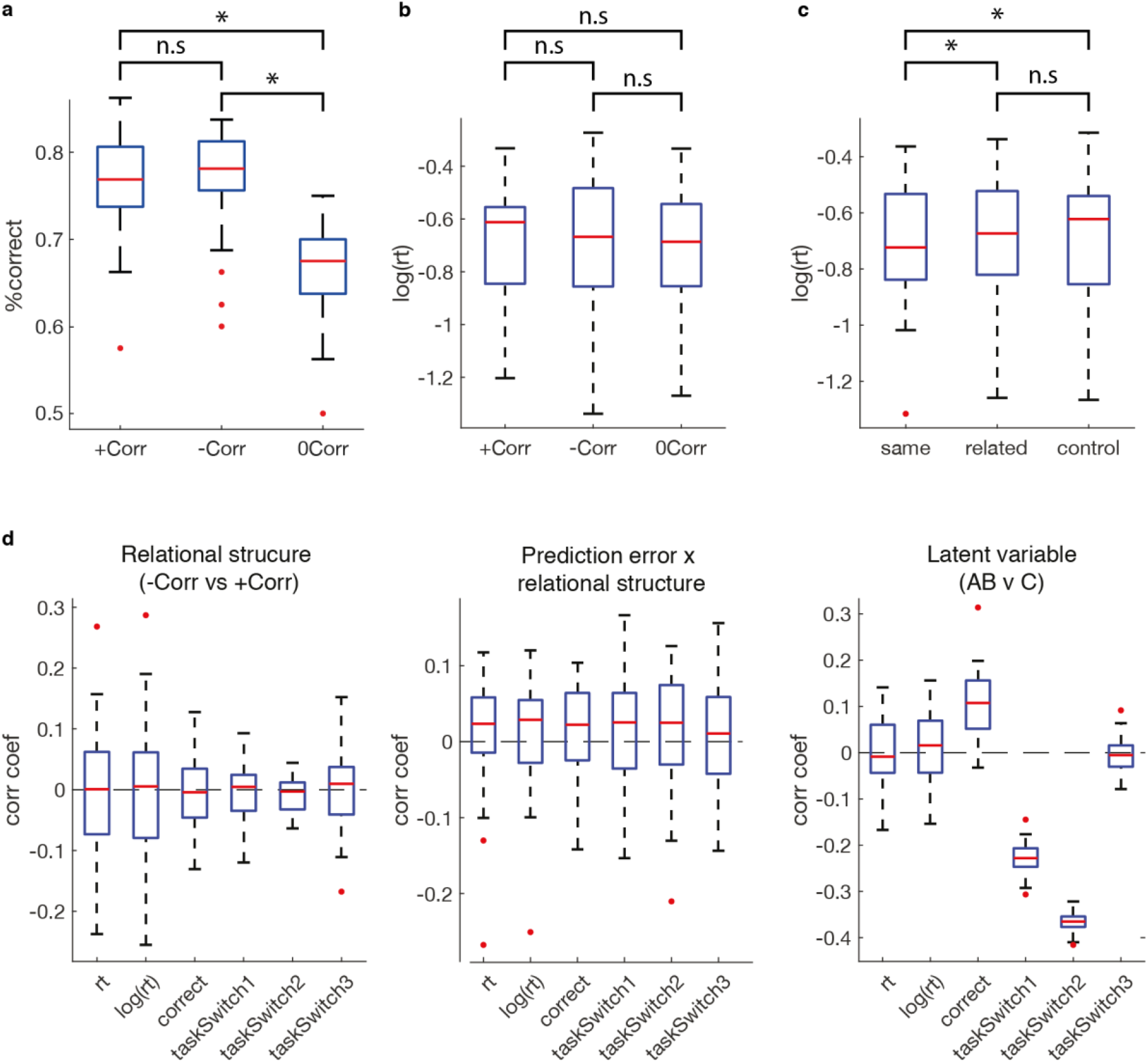
Additional behavioural analyses and possible confounds. **a.** Subjects performance (measured as % of correct choices, according to the ground truth outcome probability) was better in trials of the related stimuli than trials of the control stimulus (two-tailed paired t-tests: +Corr vs 0Corr t(27)=6.7, P<10^-6; -Corr vs 0Corr paired t(27)=7.11, P<10^-6, -Corr vs +Corr paired t(27)=0.43 P=0.66). **b.** log(reaction times), split by correlation type of trial (-Corr, +Corr or control). Reaction times did not differ between trials of stimuli under different types of correlations (two-tailed paired t-tests: P>0.4 for all comparisons). **c.** Task switching effects. We split all trials to three groups according to the relationship between current and previous trial: “same” (A→A (A trials preceded by an A trial), B→B, C→C), “related” (A→B, B→A) and “control” (A→C, B→C, C→A, C→B), and compared the means of log(reaction times) of these groups across subjects. As expected, subjects were quicker to respond to stimuli preceded by the same stimulus (one-tailed paired t-test on log(reaction times): same vs related: t(27)=2.1, P=0.02, same vs control: t(27)=2.88, P=0.003). However, there was no significant difference between log(reaction times) in trials of stimuli presented after their related stimulus (“related”) compared to trials where stimuli were presented after an unrelated stimulus (“control”), though a weak trend in the expected direction was observed (related vs control: t(27)=0.41, P=0.34). **d.** Correlation coefficients of possible behavioural confounds with the effects of interest. We first constructed six confound regressors: *reaction time, log(reaction time), correct* (−1/1 for trials where the subject’s choice was incorrect/correct according to the ground truth outcome probability, respectively), and three task switching regressors where we partitioned the three “task switching groups” (same, related, control – see panel c.) in different ways, reflecting possible levels of task switching: *taskSwitch1* (“same”: −1, “related”: 0, “control”: 1), *taskSwitch2* (“same” and “related”: −1, “control”: 1) and *taskSwitch3* (“same”: −1, “related” and “control”: 1). Next we constructed regressors reflecting our effects of interest: *relational structure* (-Corr: −1, +Corr: 1, control trials were ignored), *correctness prediction error x relational structure interaction, latent variable* (control (C) trials: −1, related (AB) trials: 1). The *relational structure* and *prediction error x structure interaction* regressors showed no significant correlations with any of the confound regressors. The *latent variable* regressor showed a positive correlation with the *correct* regressor (two-tailed paired t-test across subjects on z-transformed correlation coefficients: t(27)=7.5, P<10^-7; see panel a. for further exploration) and with *taskSwitch1* (t(27)=-35.05, P<0.0001) and *taskSwitch2* (t(27)=-84.16, P<0.0001). The latter two negative correlations were expected from the definition of the regressor (e.g. 4 out of 5 trials in [“same” + “related”] groups are either A or B trials). This is not a coincidence: A task switching effect between AB and C trials is part of the effect we were interested in when searching for a brain region that represents A and B trials more similarly than A and C or B and *C*. In all plots, the red central line is the median, the box edges are the 25^th^ and 75^th^ percentiles, the whiskers extend to the most extreme data points that are not considered outliers, and the outliers are plotted as red circles.

**Figure S3.**
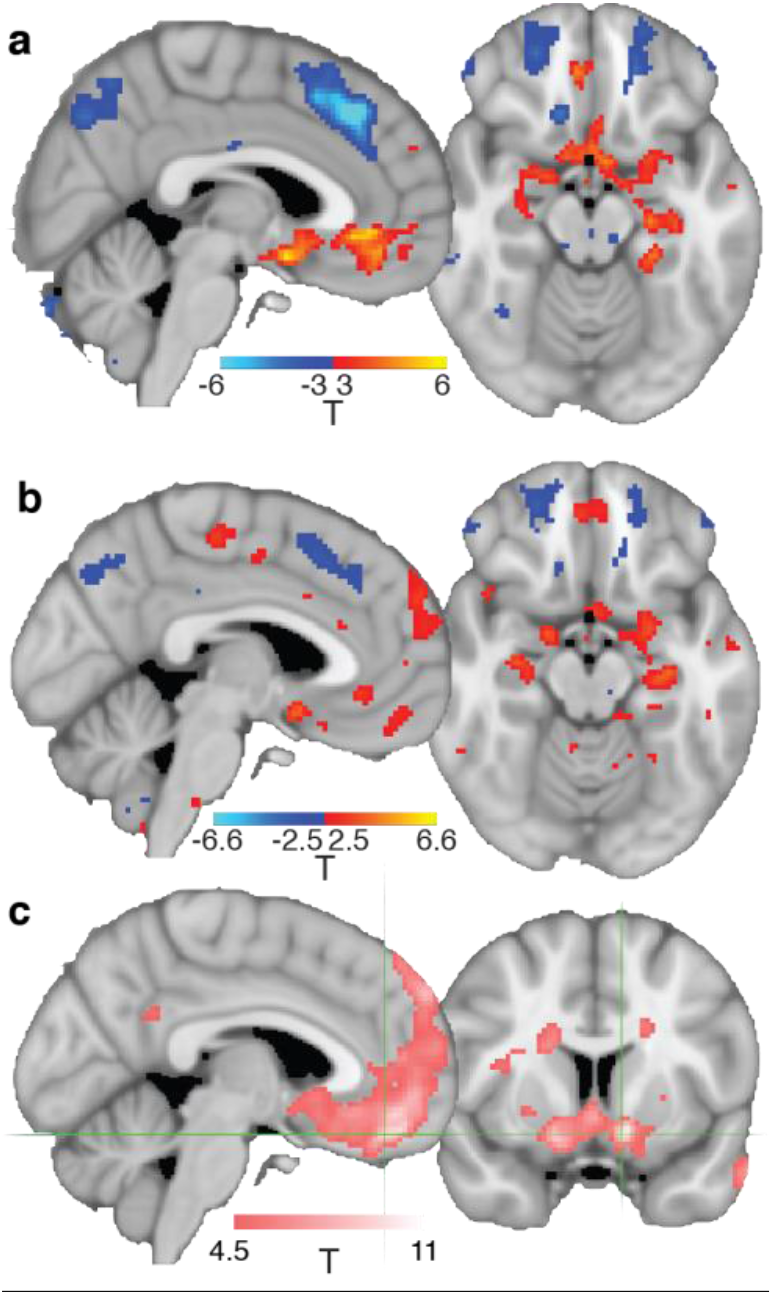
Control analyses for the univariate contrasts. **a.** Effect of the contrast [STRUCT chosen action value] > [baseline] (Same contrast as is Figure 2d), in a GLM that is similar to GLM1 but includes reaction time as a covariate. **b.** Effect of the contrast [STRUCT chosen action value] > [NAÏVE chosen action value] from the same GLM as in Figure 2d (GLM1). **c.** Effect of the contrast [STRUCT correctness prediction error] > [baseline] (same contrasts as the insets in Figure 4b and 4c), in a GLM that is similar to GLM3 but includes reaction time as a covariate.

**Figure S4.**
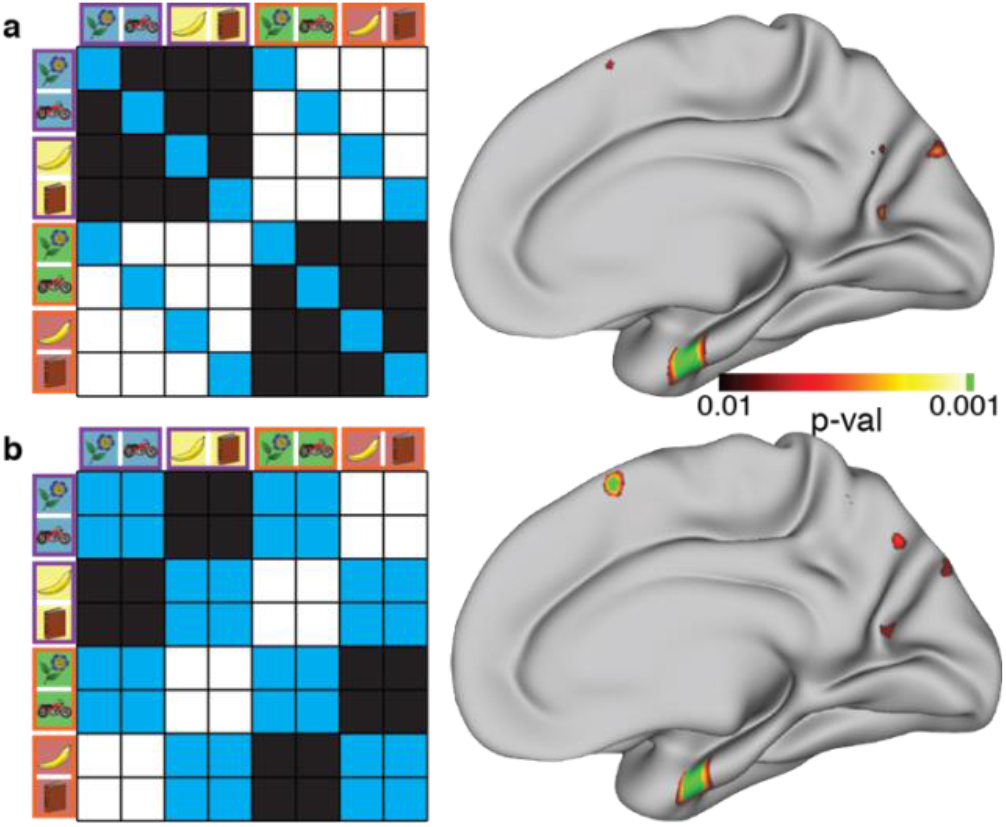
Control contrasts for the EC relational structure effect (Figure 3, top). Left: model RDMs. Black elements should be similar, white elements should be dissimilar, blue elements are ignored. Right: whole-surface results (right hemisphere). color map is the same as in **Figure 3**. **a.** Relational structure effect, ignoring all elements of pairs of same-stimulus conditions. **b.** Relational structure effect, ignoring all elements of pairs of same stimuli set conditions.

**Figure S5.**
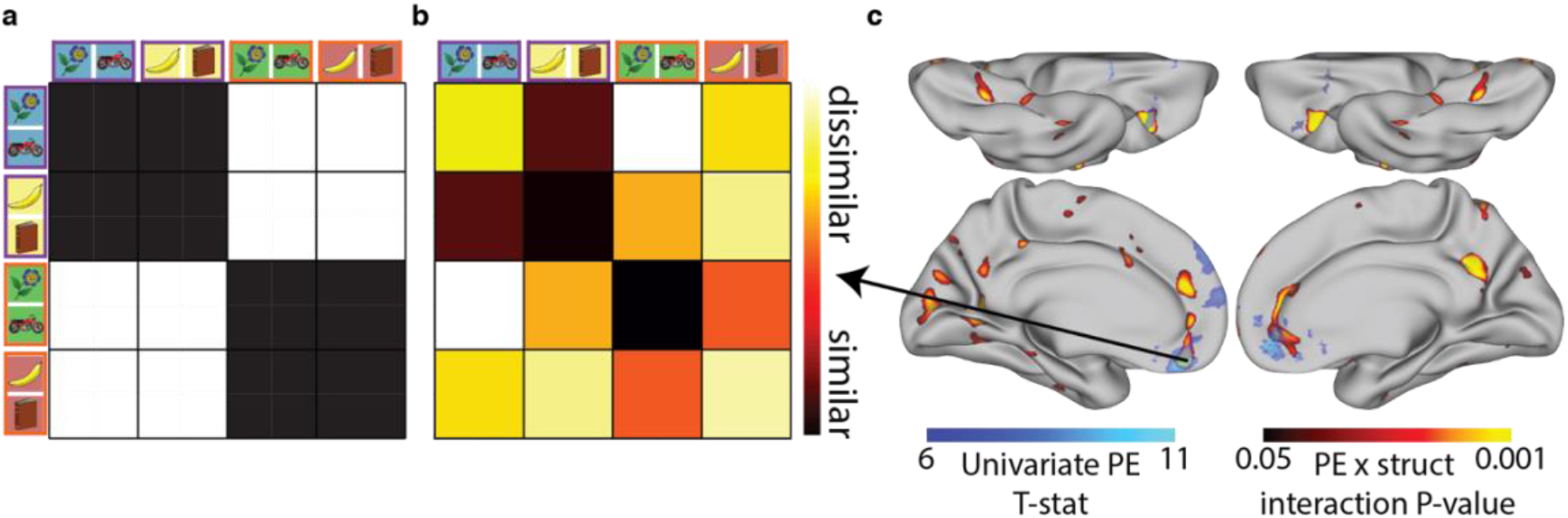
Prediction error x relational structure interaction effect. **a.** Model RDM. Note that each pair of related stimuli correspond to only one condition in the RDM as the trials of each pair are collapsed in GLM3. **b.** Visualisation of the data RDM of the prediction error x relational structure interaction effect at the peak of the vmPFC univariate prediction error effect (MNI [-4,44,-20]). **c.** Visualisation of whole-surface results of the multivariate prediction error x relational structure interaction effect (red-yellow, effects do not survive FWE-correction across a cortical hemisphere - these are exploratory results; note the low threshold), overlaid on the univariate prediction error effect, used to define the ROIs (blue – same as in the insets of Figure 4b and 4c but with a higher threshold). We found bilateral effects in vmPFC (peak P-values: LH: P=0.002 uncorrected, Figure 4a,b; RH: P=0.01 uncorrected, Figure 4c), in close proximity to each hemisphere’s univariate peak. The strongest effects in cortex where observed in PCC RH (right hemisphere): P=0.001 uncorrected) and vlPFC (LH: P < 0.001 uncorrected, P=0.002 at the peak of the univariate prediction error vlPFC effect). In all of these regions, the effects were also observed bilaterally, albeit weaker (PCC LH: P = 0.005 uncorrected; vlPFC RH: P=0.005 uncorrected). This network of regions is of particular interest for this effect, as it brings together several seemingly disparate literatures. The vmPFC and vlPFC are central to reward-guided learning [1]. The vmPFC [2–5], and PCC [4,5] have been strongly implicated in the representation of knowledge schemas. In addition, grid-like coding was reported in both vmPFC and PCC [6,7].

**Figure S6.**
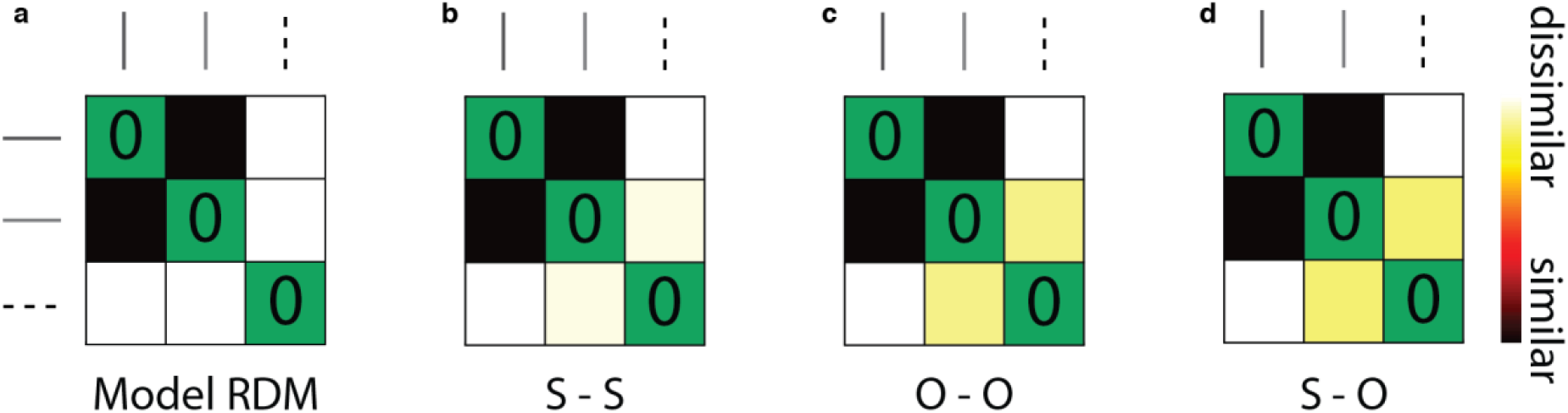
**a.** Model RDM for the latent variable within-block analysis (Figure 5). **b-d.** Visualisation of data RDMs from the latent variable effect OFC peak (MNI [-18,30,-22]), for the three different periods tested (S: stimulus presentation time; O: Outcome presentation time). The solid lines represent the two related (A and B) stimuli in each block, while the dashed line represents the control (C) stimulus (as in Figure 1b).

**Table S1.**
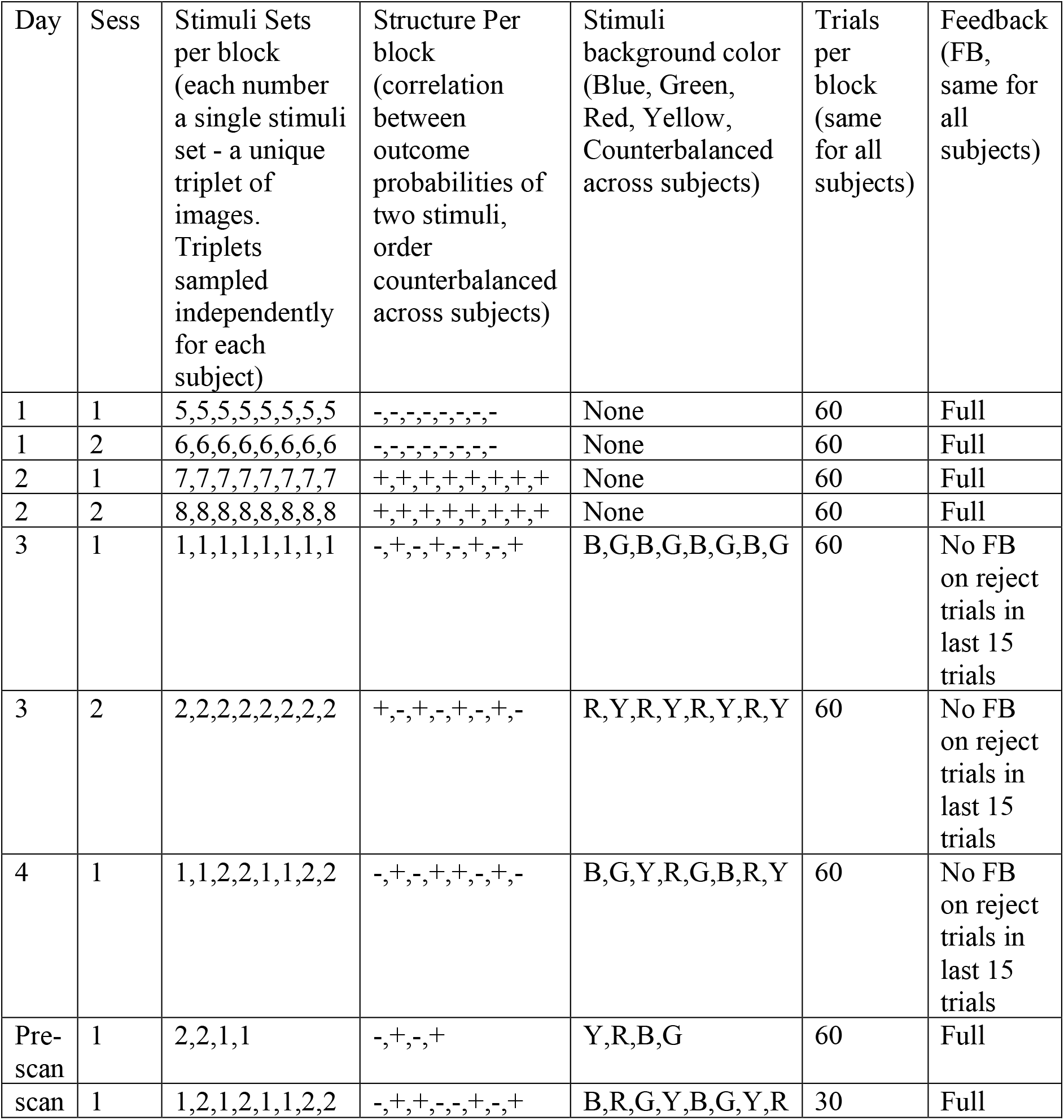
Training schedule for an example subject.

**Table S2.** Formal model comparison between STRUCT and NAÏVE models. NAIVE parameters: learning rate and inverse temperature. STRUCT parameters: learning rate, inverse temperature and 3 cross-terms.

| 2. | Model | # parameters | Log likelihood | AIC | BIC |
| --- | --- | --- | --- | --- | --- |
|  | NAÏVE | 2 params x<br>2 structures x 28 subjects = 112 | -4515.36 | 9254.72 | 10077.2 |
|  | STRUCT | 5 params x<br>2 structures x 28 subjects = 280 | -2894.44 | 6348.89 | 8405.06 |

**Table S3.**
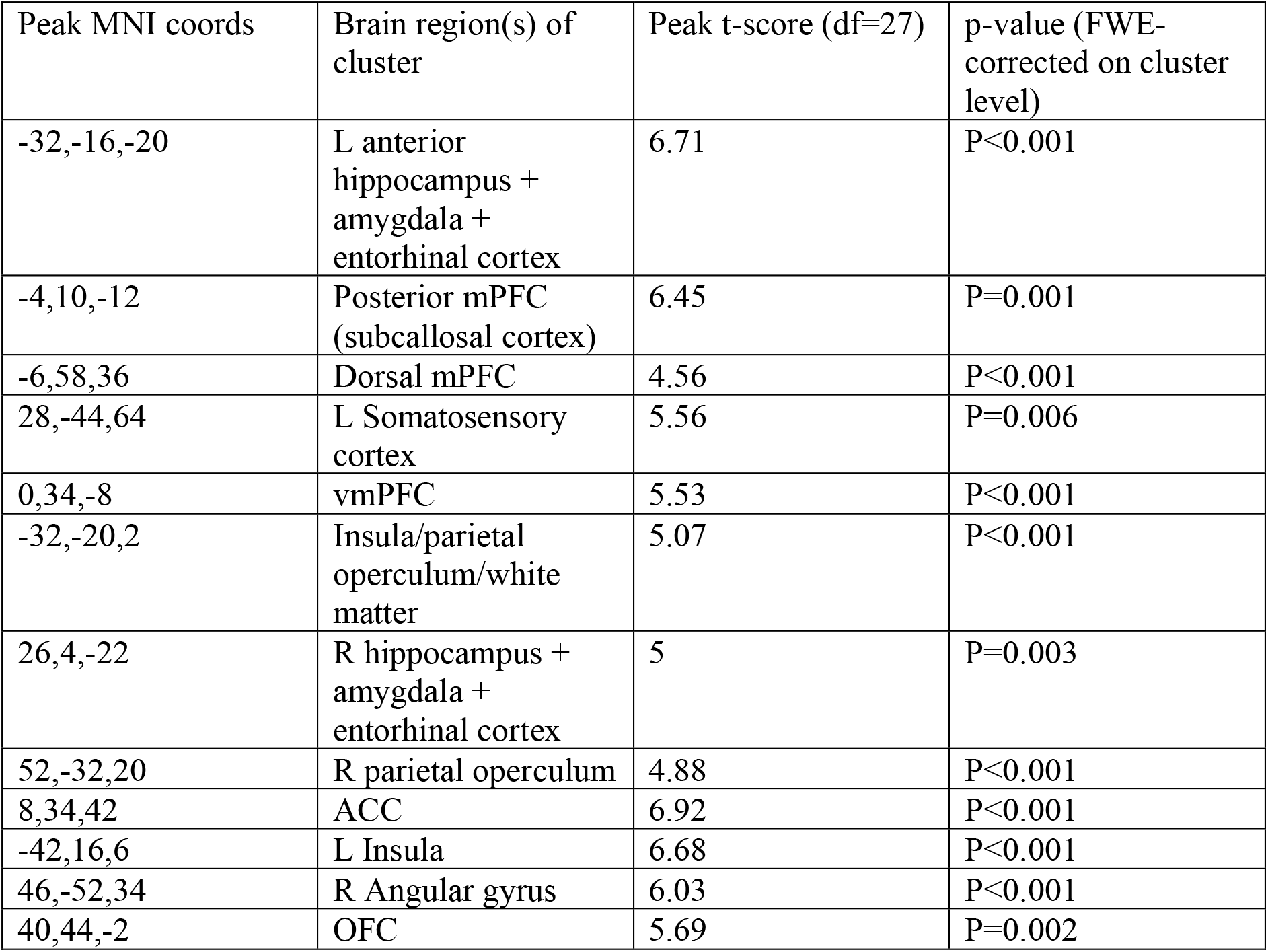
Effect of chosen action value of STRUCT model, when competing with NAÏVE model in the same GLM (GLM1, Fig. 2d). The contrast is [STRUCT chosen action value] > Baseline. All clusters with a FEW-corrected P-value < 0.05 are reported. Note that negative effects (with a negative t-score) are also reported.

| Peak MNI coords | Brain region(s) of cluster | Peak t-score (df=27) | p-value (FWE-corrected on cluster level) |
| --- | --- | --- | --- |
| -32,-16,-20 | L anterior hippocampus + amygdala + entorhinal cortex | 6.71 | P<0.001 |
| -4,10,-12 | Posterior mPFC (subcallosal cortex) | 6.45 | P=0.001 |
| -6,58,36 | Dorsal mPFC | 4.56 | P<0.001 |
| 28,-44,64 | L Somatosensory cortex | 5.56 | P=0.006 |
| 0,34,-8 | vmPFC | 5.53 | P<0.001 |
| -32,-20,2 | Insula/parietal operculum/white matter | 5.07 | P<0.001 |
| 26,4,-22 | R hippocampus + amygdala + entorhinal cortex | 5 | P=0.003 |
| 52,-32,20 | R parietal operculum | 4.88 | P<0.001 |
| 8,34,42 | ACC | 6.92 | P<0.001 |
| -42,16,6 | L Insula | 6.68 | P<0.001 |
| 46,-52,34 | R Angular gyrus | 6.03 | P<0.001 |
| 40,44,-2 | OFC | 5.69 | P=0.002 |

## Figures for shared data

25 subjects have consented to share their data, while 3 subjects did not consent.

The data of the 25 subjects who have consented can be found in https://openneuro.org/datasets/ds002801, including all the analysis levels needed to produce the figures in the paper: raw data, preprocessed data, first-level GLMs (including group-level univariate analyses), RDMs and group level RSA analyses (output of PALM). All code can be found here: https://github.com/alonbaram2/realtionalStructure_NN2020.

To aid the comparison between the paper and the shared data, in this section we provide the main figures of the paper when including data only from the subset of the 25 subjects. All main results of the paper hold within this subset (though with 25 subjects the prediction error x relational structure is significant only in LH, while RH is just below significance).

